# Membrane-anchored influenza neuraminidase vaccine drives human-like broadly protective B cell responses

**DOI:** 10.64898/2026.05.13.724804

**Authors:** Rui Liu, Chaohui Lin, Bingxin Chu, Qian Yang, Xin Wang, Changxu Chen, Ren Sun, Xudong Wu, Zeli Zhang

## Abstract

Influenza neuraminidase (NA) is a promising target for universal flu vaccines, yet eliciting potent B-cell responses against its conserved epitopes remains challenging. Here, we developed a membrane-anchored, folding-domain-free NA (mNA) that elicited superior head-specific germinal center B cell and antibody responses compared to soluble tetrameric NA. In non-human primates, mNA immunization induced cross-reactive memory B cell (MBC) responses, expanding clones with the conserved “DR” motif in HCDR3, a hallmark of human broadly reactive NA antibodies. These MBCs conferred cross-inhibitory activity against diverse NA variants and *in vivo* cross-protection. Cryo-EM analysis revealed that the 554-C2 clone targets the conserved enzymatic pocket via the “DR” motif, while the 554-C1 clone recognizes previously uncharacterized epitopes at the interface between two adjacent N2 monomers, effectively reducing plaque formation by contemporary H3N2 strains. Our findings highlight the immunological advantages of membrane-anchoring, providing a robust strategy for designing next-generation vaccines against influenza and other pathogens.

## Introduction

Influenza infections represent a significant global public health challenge, with seasonal epidemics resulting in an estimated 250,000 to 650,000 deaths annually^1,2^. Current seasonal influenza vaccines are primarily based on the virus’s hemagglutinin (HA) protein and are designed to target specific strains predicted to circulate in the upcoming season^3^. However, these licensed vaccines offer suboptimal efficacy, with protection rates varying between 10% and 60% against seasonal strains and provide limited defense against emerging pandemic influenza strains^4–8^. Consequently, there is a global imperative to develop a universal influenza vaccine that confers broad and durable protection across diverse viral strains^9^.

In contrast to HA, NA evolves more slowly and harbors a greater number of conserved epitopes, making it a promising target for universal influenza vaccines^10,11^. Growing evidence highlights the critical role of NA-specific immunity in reducing disease severity and limiting viral shedding in infected individuals^12–16^. Notably, several broadly inhibitory monoclonal antibodies (BImAbs) have been identified in humans that target highly conserved enzymatic epitopes on the NA head^17–21^. These NA BImAbs, often characterized by a long heavy-chain complementarity-determining region 3 (HCDR3) containing a “DR” motif, demonstrate broad cross-reactivity across diverse influenza NA subtypes and confer robust protection *in vivo*^22^, providing valuable insights for the development of NA-based universal vaccines. Although several NA vaccine candidates that utilize exogenous folding domains to tetramerize the NA head have been evaluated in murine models^23–26^, the immune competition and off-target effects of these exogenous folding domains remain unresolved. In addition, no NA vaccines have been tested in non-human primates (NHPs) for their capacity to elicit broadly reactive B cell responses.

Here, we demonstrated that membrane-anchored, folding-domain-free NA (mNA) elicits stronger head-directed and functional humoral responses than exogenous folding-domain-tetramerized soluble NA (sNA). Notably, immunization of rhesus macaques with mNA induced NA cross-reactive memory B cells expressing the “DR” motif in the HCDR3, a hallmark of NA BImAbs identified in humans^22^. These findings underscore the enhanced immunogenicity of mNA and its capacity to expose conserved, functionally relevant epitopes on the NA head domain, thereby promoting cross-reactive B cell responses.

## Results

### The mNA immunization recruited superior NA head-specific precursor B cells and induced stronger functional antibody responses

The tetramerization of NA is essential for its immunogenicity^27–29^, and several exogenous folding domains have been utilized to tetramerize the NA head^20,30^. To overcome the immunogenicity and potential immune interference associated with exogenous folding domains, we developed mN2, a full-length immunogen comprising the head, stalk, transmembrane domains, and cytoplasmic tail, which resembles the native NA sequences on the influenza virion (Figure 1A). Both sN2 and mN2 were efficiently expressed in insect cells (Figure 1B). Following tag-affinity purification, sN2 formed higher-order multimers, whereas mN2 is more homogeneous and predominantly assembled into tetramers (Figures S1A-S1C). When formulated in a solution containing 0.02% polysorbate 80 (PS80), mN2 underwent multimerization, with multiple NA heads displayed on detergent micelles (Figure S1D).

**Figure 1.**
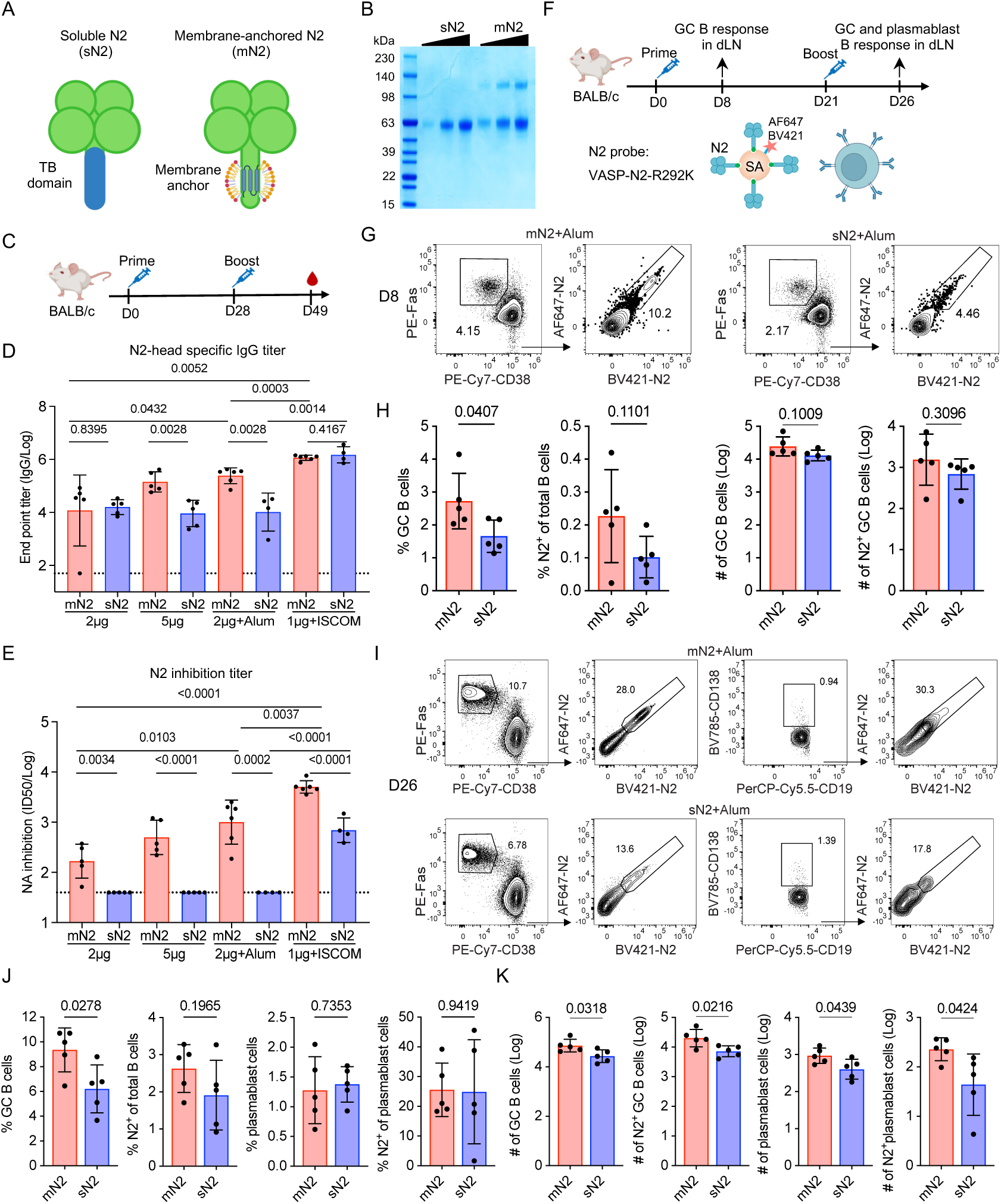
The mN2 immunization recruited superior N2 head-specific precursor B cells in GC and elicited higher functional antibody titers. **(A)** Schematic illustration of sNA and mNA. TB stands for the tetramerization domain derived from *Staphylothermus marinus* tetrabrachion. **(B)** Purified sN2 and mN2 immunogens (A/Kansas/14/2017-H3N2) were analyzed by SDS-PAGE and Coomassie blue staining. **(C to E)** Female mice (n = 4-6 per group) were prime-boosted at the indicated time and dose with sN2 or mN2. One adjuvant group received 2 μg sN2 or mN2 with 50 μg Alum adjuvant; the other adjuvant group received 1 μg sN2 or mN2 with 5 μg ISCOM adjuvant. N2 head-specific IgG antibody levels in sera were measured by ELISA using VASP-N2-coated plates **(D).** The anti-N2 NAI titers were determined by ELLA (E). **(F to K)** Female BALB/c mice (n = 5 per group) were prime-boosted at the indicated time and dose with 62.5 µg Alum adjuvant combined with either 2.5 µg of mNA or sNA. The GC B and plasmablast B responses in the draining lymph nodes were analyzed on day 8 and day 26. N2-specific B cells were identified by the mutant N2 head-specific probe **(F)**. Representative gating strategy for GC B cells (live^+^CD19^+^Fas^+^CD38^-^), N2-specific GC B cells (live^+^CD19^+^Fas^+^CD38^-^N2-AF647^+^N2-BV421^+^), plasmablast B cells (live^+^CD19^+^IgD^-^CD138^+^), and N2-specific plasmablast B cells (live^+^CD19^+^IgD^-^CD138^+^N2-AF647^+^N2-BV421^+^) on day 8 (G) and day 26 (I), respectively. The frequencies and cell numbers of GC B cells and N2-specific B cells (relative to total B cells) on day 8 are compared (H). The frequencies (J) and cell numbers (K) of GC B cells, N2-specific B cells (relative to total B cells), plasmablast B cells, and N2-specific plasmablast B cells on day 26 are compared. Raw data (H-left panel and J) or log-transformed data (D, E, H-right panel, and K) were presented as mean ± s. d. Statistical analyses were performed with raw or log-transformed data using the two-tailed unpaired *t*-test, and *p*-values were indicated.

To evaluate the immunogenicity of sN2 and mN2, eight groups of mice were prime-boosted with varying doses of the indicated immunogens, formulated with or without Alum or saponin-based nanoparticle (ISCOM) adjuvants (Figures 1C-1E). We found that the mN2 elicited significantly higher levels of head-specific and anti-N2 inhibition titers than sN2, with responses showing dose dependency (Figures 1D and 1E). Including Alum adjuvant further boosted the N2-inhibiting antibody titers but did not improve the functional inhibitory antibodies elicited by sN2. ISCOM adjuvant significantly increased inhibitory antibody titers for both antigens, with the mN2 group achieving titers approximately 10-fold higher than those induced by sN2 (Figure 1E). Mechanistically, we observed that a single dose of mN2 elicited a significantly higher frequency of GC B cells in the draining lymph nodes (dLNs) compared to a single dose of sN2 (Figures 1F–1H), and the frequency of N2 head-specific GC B cells among total B cells in the dLNs showed an increasing trend in the mN2 group, while the absolute numbers were similar between the mN2 and sN2 groups (Figure 1F-1H). Notably, after boosting, the absolute numbers of both N2 head-specific GC B cells and plasmablasts were significantly elevated in the mN2-immunized group (Figures 1I–1K), indicating that mN2 more effectively recruited head-specific B cell precursors and mitigated immune competition from the TB-folding domain. Moreover, mN2 combined with ISCOM generated superior inhibitory antibody responses via both subcutaneous and intramuscular routes compared to sN2 with ISCOM (Figures S2A-S2D). To exclude potential confounding effects of detergent in the formulation, an equivalent concentration of PS80 was added to sN2 prior to immunization; however, this did not improve the weak inhibitory antibody response (Figures S2E-S2G).

We also expressed mN1 and sN1 in insect cells (Figure 2A). Immunization with mN1 elicited significantly higher N1 head-specific IgG and anti-N1 inhibition titers, whereas no inhibitory antibodies were detected in the sN1 group (Figures 2B-2D). Consistently, immunization with mN1 conferred improved protection against the antigenically distinct A/California/04/2009-H1N1 than sN1 immunization (Figures 2E). Collectively, these findings demonstrate that mNA is superior to sNA in eliciting head-specific and functional antibody responses, and that immunization with mNA provides protection against influenza infection.

**Figure 2.**
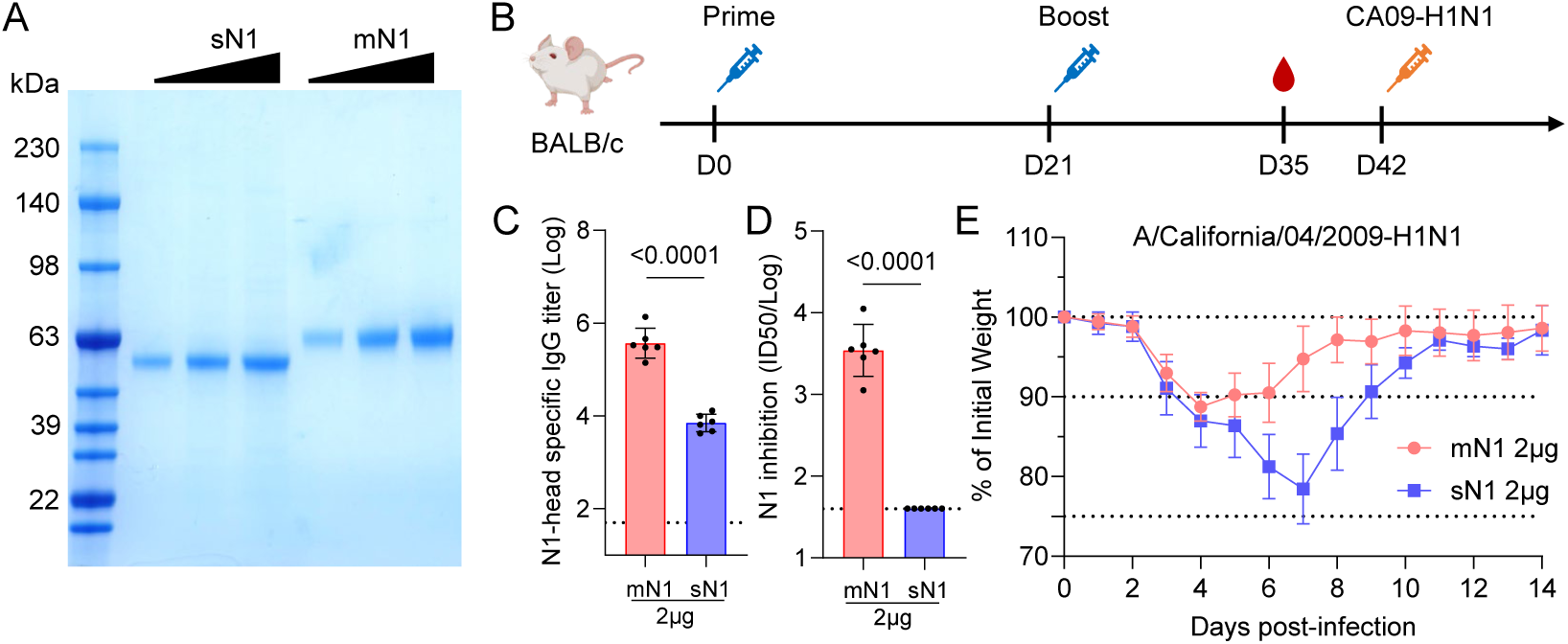
The mN1 immunization provides greater protection than sN1. **(A)** Purified sN1 and mN1 immunogens (A/Michigan/45/2015-H1N1) were analyzed by SDS-PAGE and Coomassie blue staining. **(B-E)** Female mice (n = 6 per group) were prime-boosted at the indicated time and dose with sN1 or mN1, and then challenged with ∼5xmLD50 CA09-H1N1. N1 head-specific IgG antibody levels in sera were measured by ELISA using VASP-N1-coated plates (C). The anti-N1 NAI titers were determined by ELLA (D). The percentage of body weight after CA09-H1N1 challenge was shown (E). The data are presented as mean ± s. d. Statistical analyses were performed with log-transformed data using the two-tailed unpaired *t*-test (C and D), and *p*-values were indicated.

### Vaccinating the NHPs with mNA generated cross-reactive inhibiting antibody responses

Previous studies have reported pre-existing humoral immunity to N1 and N2 in healthy individuals^20,31^, which likely modulates the immune response induced by subsequent vaccination^32^. To assess the immunogenicity of the mN1+mN2 vaccine in the context of pre-existing NA immunity, we established such immunity in mice through sequential immunization with mN1 and mN2 (Figure S3A). Initial immunization with mN1 induced N1 head-specific antibody responses with limited cross-reactivity to N2. Subsequent immunization with mN2 boosted anti-N2 antibody responses and enhanced N1-inhibiting antibody titers, indicative of a cross-reactive memory B cell response (Figure S3B-S3E). Under conditions of pre-existing NA immunity, administration of the mN1+mN2 vaccine further increased both N1- and N2-specific head-directed binding antibodies and NI titers. Notably, at the final time point, NI titers against both N1 and N2 were significantly higher in the ISCOM-adjuvanted group compared to the Alum group. These findings demonstrate that the mN1+mN2 vaccine retains robust immunogenicity despite pre-existing NA immunity.

We next evaluated the immunogenicity of mNA in rhesus macaques, which were assigned to two groups receiving either Alum or ISCOM as adjuvant (Figure 3A). Given Alum’s lower adjuvant potency, the immunogen dose in this group was doubled relative to the ISCOM group. Pre-existing NA immunity was first established by administering mN1 on day 0 and mN2 on day 21, followed by two booster immunizations with the mN1+mN2 vaccine. Blood samples were collected on days 0, 21, 42, 63, 77, 91, 109, and 330 to assess the durability of NA-specific antibody responses (Figure 3A). No substantial differences in NA-specific antibody responses were observed between the Alum and ISCOM groups (Figures 3B-3E). As observed in mice, priming with mN1 elicited N1 head-specific antibodies with minimal cross-reactivity to N2. Subsequent mN2 immunization boosted IgG titers against both N1 and N2, indicating cross-reactive antibody responses. Following the establishment of pre-existing immunity, the first mN1+mN2 vaccination rapidly enhanced both N1- and N2-specific binding antibodies. The second booster further increased IgG titers, which peaked on day 91 and persisted up to day 330, albeit at reduced levels. The decline from peak titers was approximately 32-fold (Alum) and 13-fold (ISCOM) for N1, and 100-fold (Alum) and 27-fold (ISCOM) for N2. Notably, the ISCOM group exhibited smaller fold-reductions, suggesting improved antibody persistence (Figures 3B and 3C).

**Figure 3.**
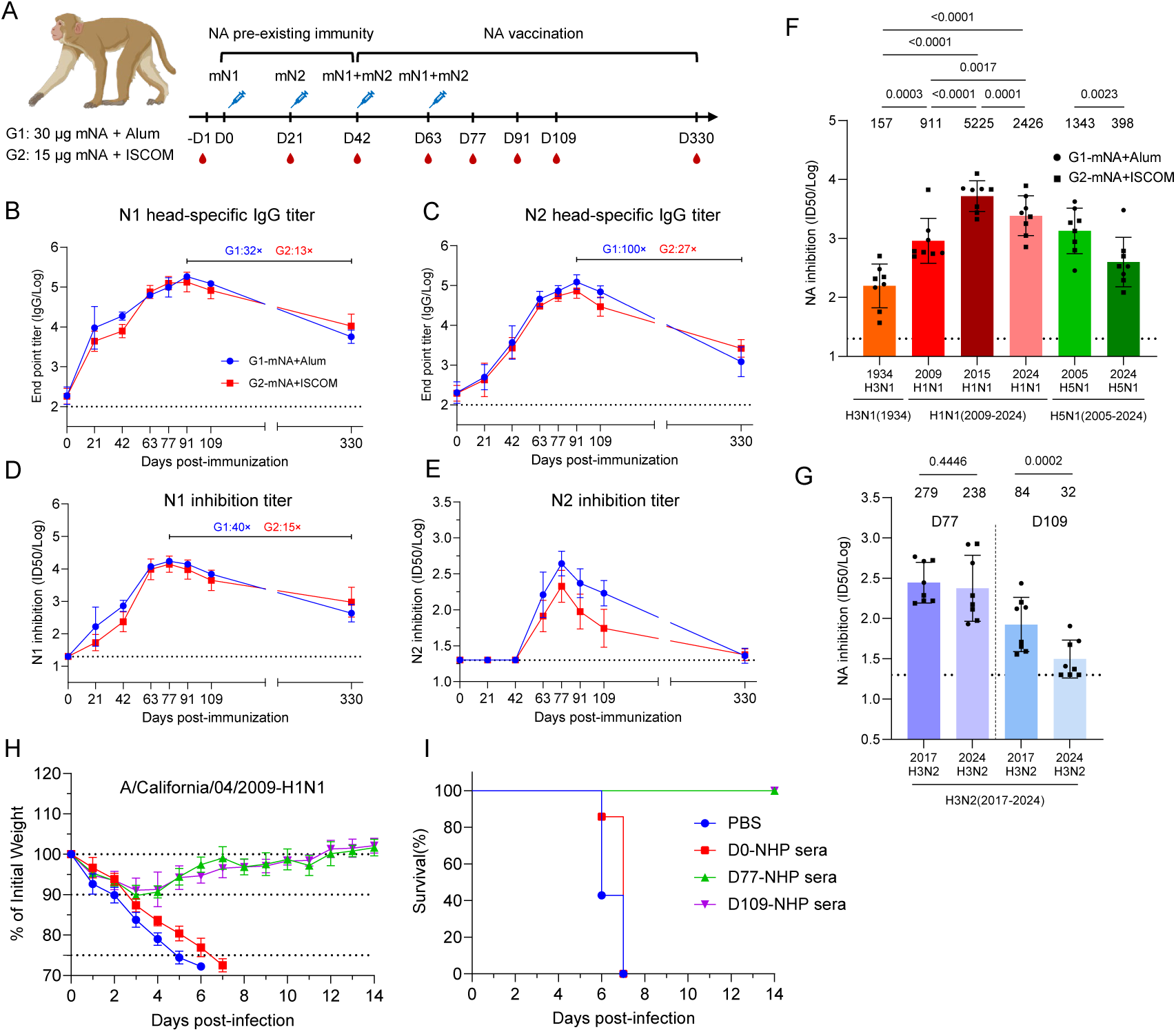
mNA elicited cross-reactive and protective antibody responses in NHPs. **(A-E)** Rhesus macaques (n = 8) were immunized with mNA at the indicated dose and time. Group 1 (G1, n = 3, blue) received 30 µg mNA with Alum adjuvant. Group 2 (G2, n = 5, red) received 15 µg mNA with ISCOM adjuvant (A). N1 (B) or N2 (C) head-specific IgG levels in NHP sera were measured by ELISA. N1 (D) or N2 (E) NAI titers were assessed by ELLA. Fold changes in antibody titers between peak and day 330 were shown (B-D). **(F and G)** The NAI titers against indicated NAs derived from N1, H5N1, or N2 were detected by ELLA. The geometric means of the NAI titers are shown in each colum. Data were log-transformed and presented as mean ± s. d. Statistical analyses were performed with log-transformed data using a paired *t*-test, and *p*-values were indicated. (**H and I**) The percentage of initial body weight and survival rates in BALB/c mice following passive transfer of the indicated NHP immune sera (eight NHP sera were equally pooled). Mice (n = 6 per group) received an intraperitoneal injection of the indicated NHP immune sera or PBS. 12 hours post sera-transferring, mice were challenged with ∼5 × mLD_50_ of A/California/04/2009 (H1N1). The percentage of body weight and the survival rate were shown. The data are presented as mean ± s. d.

The kinetics of N1- and N2-specific NI antibody responses mirrored those of binding antibodies (Figures 3D and 3E). Both N1- and N2-specific NI titers peaked on day 77 and gradually declined thereafter. N1-specific NI titers remained detectable on day 330, with 40-fold (Alum) and 15-fold (ISCOM) reductions from peak levels (Figure 3D). In contrast, N2-specific NI titers declined more substantially, reaching the limit of detection by day 330. Comparable magnitudes and kinetics of N2-specific NI responses were observed using protein- and live virus-based assays (Figures S4A and S4B). This accelerated waning of N2-specific immunity aligns with epidemiological data showing faster decline in vaccine effectiveness against H3N2 compared to H1N1^33^, underscoring the need for improved H3N2 vaccine formulations to achieve durable protection^6^.

We further examined cross-reactive NI responses against diverse NA subtypes. Sera collected on day 109 exhibited inhibitory activity against N1 from 1934-H3N1 (PR8, H3-1968), 2009-H1N1, 2015-H1N1, 2024-H1N1, 2005-H5N1, and 2024-H5N1, with the highest titers directed against the homologous 2015-H1N1 strain (Figure 3F). Cross-inhibitory antibodies against N2 from 2017-H3N2 and 2024-H3N2 were also detected in sera of day 77 and day 109 (Figure 3G). These results indicate that the mN1+mN2 immunization elicits broad and cross-reactive NA-specific antibody responses in NHPs.

### Passive serum transfer from mNA-vaccinated NHPs confers protection against antigenically distinct H1N1 challenge in mice

To evaluate the protective efficacy of antibodies induced by mNA immunization, serum was collected from NHPs at days 0, 77, and 109 post-vaccination, equally pooled, and transferred (400 µL per mouse) to naïve mice prior to challenge with antigenically distinct H1N1 (A/California/04/2009 and A/Jiangsu-Tinghu/SWL1586/2024). In the A/California/04/2009-H1N1 challenge model, mice that received NHP serum (day 77 and day 109 post-immunization) showed rapid recovery of body weight and achieved 100% survival, whereas all mice died in the control group (Figures 3H and 3I). The pathogenicity of A/Jiangsu-Tinghu/SWL1586/2024-H1N1 was lower than that of A/California/04/2009-H1N1 in mice, and the transferred day 77 and day 109 sera reduced the mortality and morbidity caused by this strain (Figures S4C and S4D). These findings demonstrate that mNA vaccination elicits functional, protective antibodies against antigenically distinct H1N1.

### The mNA vaccine elicited cross-reactive memory B cell responses and drove clonal expansion in NHPs

To evaluate NA-specific B cell responses, we employed NA mutant probes previously developed by our group^20,22^, in which key sialic acid-binding residues in N1 and N2 were mutated to minimize background staining and enable specific detection of NA-specific and cross-reactive MBCs (Figure 4A). The gating strategy for identifying N1-specific, N2-specific, N1-N2 cross-reactive, and total NA-specific MBCs is detailed in Figures S5A-S5I. At baseline (day 0), NA-specific MBCs were nearly undetectable in all immunized NHPs (Figure 4B). No significant difference in NA-specific MBC frequencies was observed between the Alum and ISCOM adjuvant groups; therefore, data from both groups were combined for subsequent analyses. By day 77 post-immunization, significant increases were observed in the frequencies of N1-specific (mean: 0.4768%), N2-specific (mean: 0.4771%), and total NA-specific MBCs (mean: 1.0313%) among IgD⁻IgG⁻^/^⁺ B cells (Figure 4B). Notably, cross-reactive NA-specific MBCs were also induced, reaching a frequency of 0.023% at day 77 (Figure 4B). These responses persisted up to day 330, albeit at reduced levels (Figures 4C, S5I, and S5J). These results demonstrate that mNA is highly immunogenic in NHPs, eliciting both strain-specific and cross-reactive MBCs.

**Figure 4.**
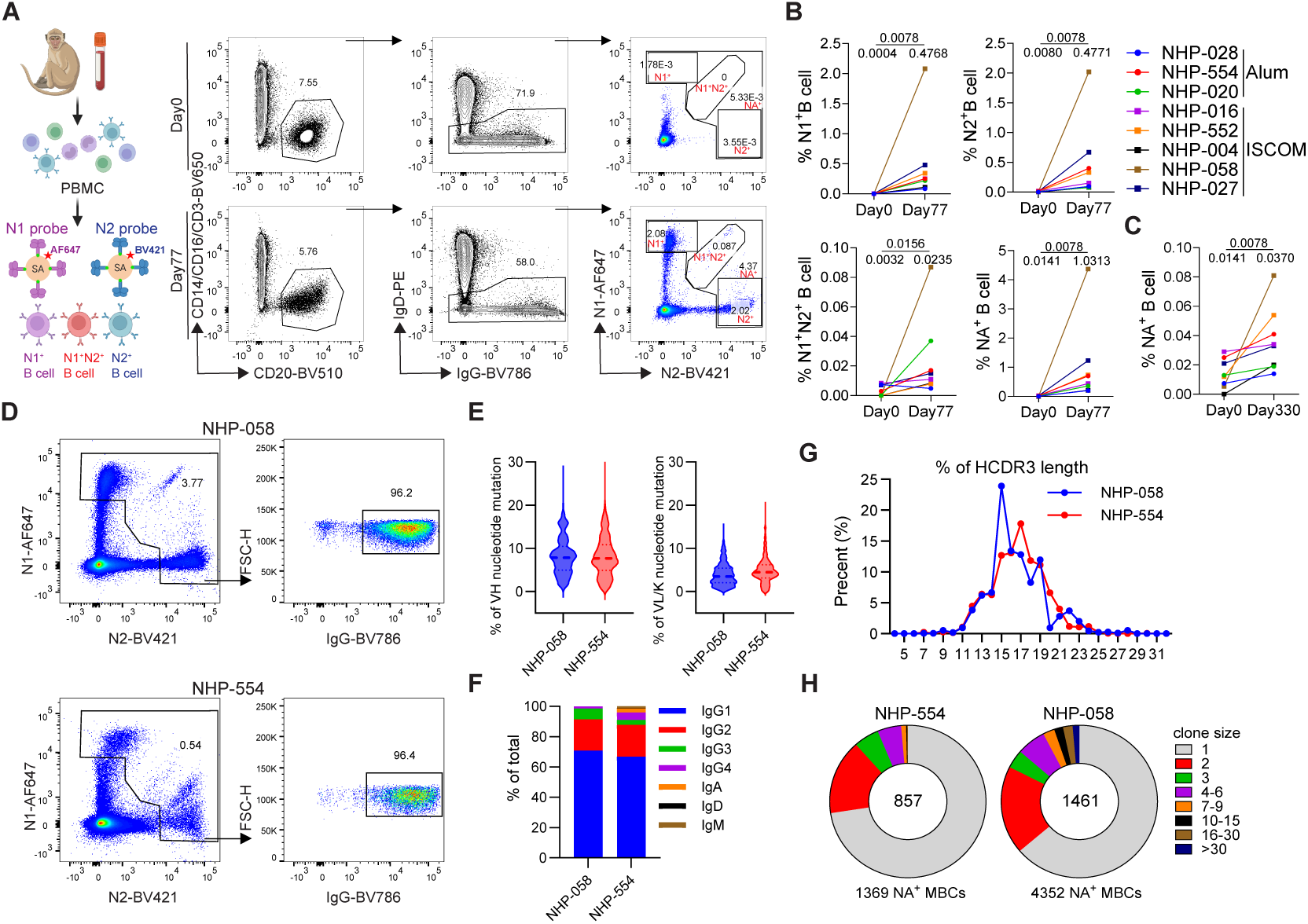
mNA elicited strain-specific and cross-reactive memory B cell responses in NHPs. **(A)** Gating strategy to identify N1-specific, N2-specific, or NA cross-reactive MBCs at day 0 and day 77. Cells were gated on live, CD14^-^, CD16^-^, CD3^-^, CD20^+^, IgD^-^, IgG^+/-^ B cells. **(B)** The frequency of N1^+^, N2^+^, N1^+^N2^+^, or NA^+^ MBCs at day 0 and day 77 was shown and compared. **(C)** The frequency of NA^+^ MBCs at day 0 and day 330 was shown and compared. The mean frequencies were indicated at each time point of each graph (B and C). **(D)** Representative flow plots show IgG^+^ B cells within sorted NA-specific MBCs in NHP-058 and NHP-554 at day 77. **(E)** VH nucleotide mutation rates of sequenced NA-specific MBCs from NHP-058 and NHP-554. **(F)** The frequency of different Ig subtypes of NA-specific MBCs from NHP-058 and NHP-554. **(G)** HCDR3 length distribution of sequenced NA-specific MBCs. **(H)** The proportion of each NA-specific MBC clonal family in NHP-058 and NHP-554. Numbers within the donut plot indicate the total number of clonal families detected for each NHP. Data in (B) and (C) were analyzed for statistical significance using the Wilcoxon matched-pairs signed-rank test, and *p*-values were indicated.

To further characterize the NA-specific MBC repertoire, we performed single-cell B cell receptor (BCR) sequencing on two NHPs (NHP-058 and NHP-554). NA-specific MBCs were isolated from CD3⁻CD14⁻CD16⁻CD20⁺IgD⁻IgG⁻^/^⁺ B cells using dual N1 and N2 mutant probes. Of the sorted cells, 96% were IgG-positive (Figure 4D), indicating robust class-switching following immunization. Both heavy and light-chain variable (V) genes exhibited high levels of somatic hypermutation (SHM) (Figure 4E). IgG1 was the predominant isotype among NA-specific MBCs, accounting for approximately 60% of sequences, followed by IgG2 at 20% (Figure 4F). Previous studies have shown that NA BImAbs typically possess long HCDR3 loops that penetrate the conserved enzymatic site of NA^22,34^. Accordingly, we analyzed the HCDR3 length distribution and found that in NHP-554, NA-specific MBCs were predominantly characterized by a 17-amino-acid HCDR3 (frequency: ∼17%), followed by 15- and 19-amino-acid HCDR3s (∼12% each) (Figure 4G). In contrast, NHP-058 exhibited a broader distribution with three dominant peaks at 15, 19, and 22 amino acids (∼25%, 12%, and 5%, respectively). Clonal diversity analysis revealed that 1,369 paired BCR sequences from NHP-554 clustered into 857 distinct clones, while 4,352 sequences from NHP-058 grouped into 1,461 clones (Figure 4H). NHP-058 displayed a higher proportion of large clones (size >7) compared to NHP-554. Together, these findings indicate that mNA vaccination induces a diverse, clonally expanded, and highly somatically hypermutated NA-specific memory B cell response in NHPs.

### The NA cross-reactive inhibiting mAbs were identified from NA-specific MBCs in NHPs

To characterize the clonal evolution of NA-specific MBCs, we analyzed the BCR sequences of the two most prevalent clonotypes from NHP-554 that harbored the “DR” motif in the HCDR3 (Figure 5A), a signature feature previously linked to broadly inhibitory anti-NA mAbs isolated from humans^22,34^. These clonotypes displayed evidence of clonal evolution, with variable levels of SHM relative to their inferred germline sequences (Figures 5B and 5C). To evaluate their functional properties, we synthesized and expressed the mAbs 554-C1-1 and 554-C2-1. 554-C1-1 demonstrated inhibitory activity against N2-2017 and N2-2024 in the ELLA, but no inhibition was observed against N2-2017 in the MUNANA assay, suggesting that 554-C1-1 likely inhibits NA through steric hindrance (Figures 5D–5G). In contrast, 554-C2-1 exhibited inhibition against diverse N1 subtypes in both ELLA and MUNANA assays, indicating targeting of the N1 enzymatic active site (Figures 5H–5P).

**Figure 5.**
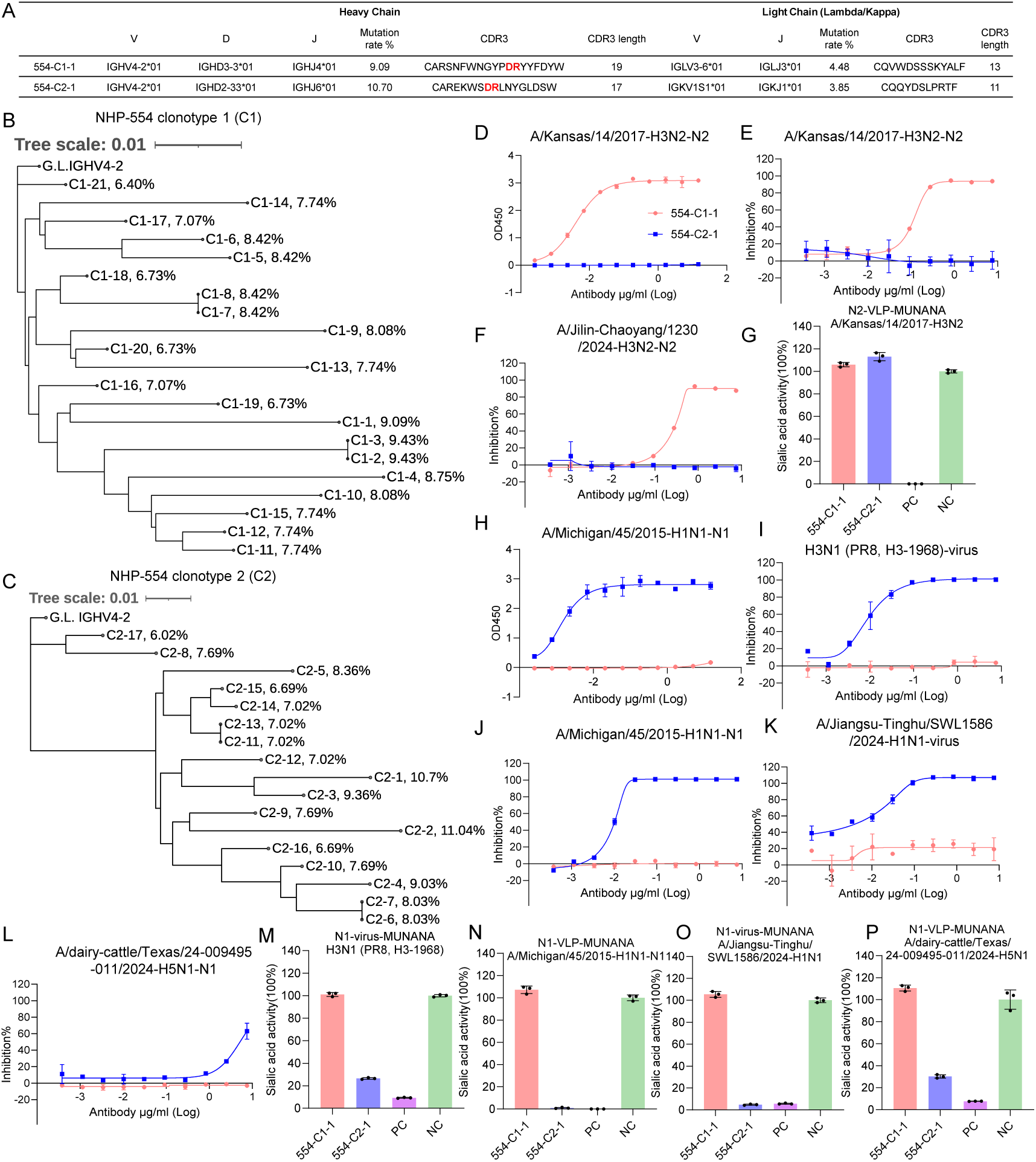
mNA induced the expansion of NA-specific B cell clones and promoted somatic hypermutation. **(A)** The VDJ usage, CDR3 lengths, and V gene mutation rates in the heavy and light chains of the two most prevalent clonotypes (mAbs 554-C1-1 and 554-C2-1) from NHP-554 were shown. **(B and C)** Clonal evolutionary trees based on heavy chain VDJ sequences for clonotype 1 (C1) and clonotype 2 (C2) from NHP-554 were shown in (B) and (C), respectively, along with their heavy chain V gene mutation rates. The scale bar represents a 0.01% change in nucleotides. **(D-P)** Binding and inhibitory activities of two NA mAbs against both N2 and N1 subtypes. The binding activity of mAbs to recombinant N2 (rN2, A/Kansas/14/2017-H3N2) (D) and N1 (rN1, A/Michigan/45/2015-H1N1) (H) was assessed by ELISA. The inhibition activities of mAbs were evaluated using ELLA against various N2 (E and F) and N1 (I to L) influenza virus strains. The data are presented as mean ± s. d. The technical duplicates were utilized for each mAb (D-F and H-L). The inhibition of NA enzymatic activity by mAbs was assessed by MUNANA assay using the indicated various influenza viruses or NA-VLPs. The technical triplicates were utilized for each mAb (G and M-P).

Subsequently, we expressed mAbs derived from the most abundant clonotypes (C3, C5, and C6 in NHP-554 and C1-C5 in NHP-058) with the highest VH mutation frequencies. Among these, only 554-C6 contained the “DR” motif in the central region of the HCDR3 (Table S1), whereas the remaining clonotypes lacked this motif. Additional mAbs derived from clonotypes harboring long HCDR3 regions with the “DR” motif (Figures 6A-6C, indicated in red, and Table S1) were also expressed; these BCRs were each represented by a single cell within the NA-specific repertoire. The V, diversity (D), and joining (J) gene usage, V gene somatic mutation rates, and CDR3 sequences are summarized in Table S1.

**Figure 6.**
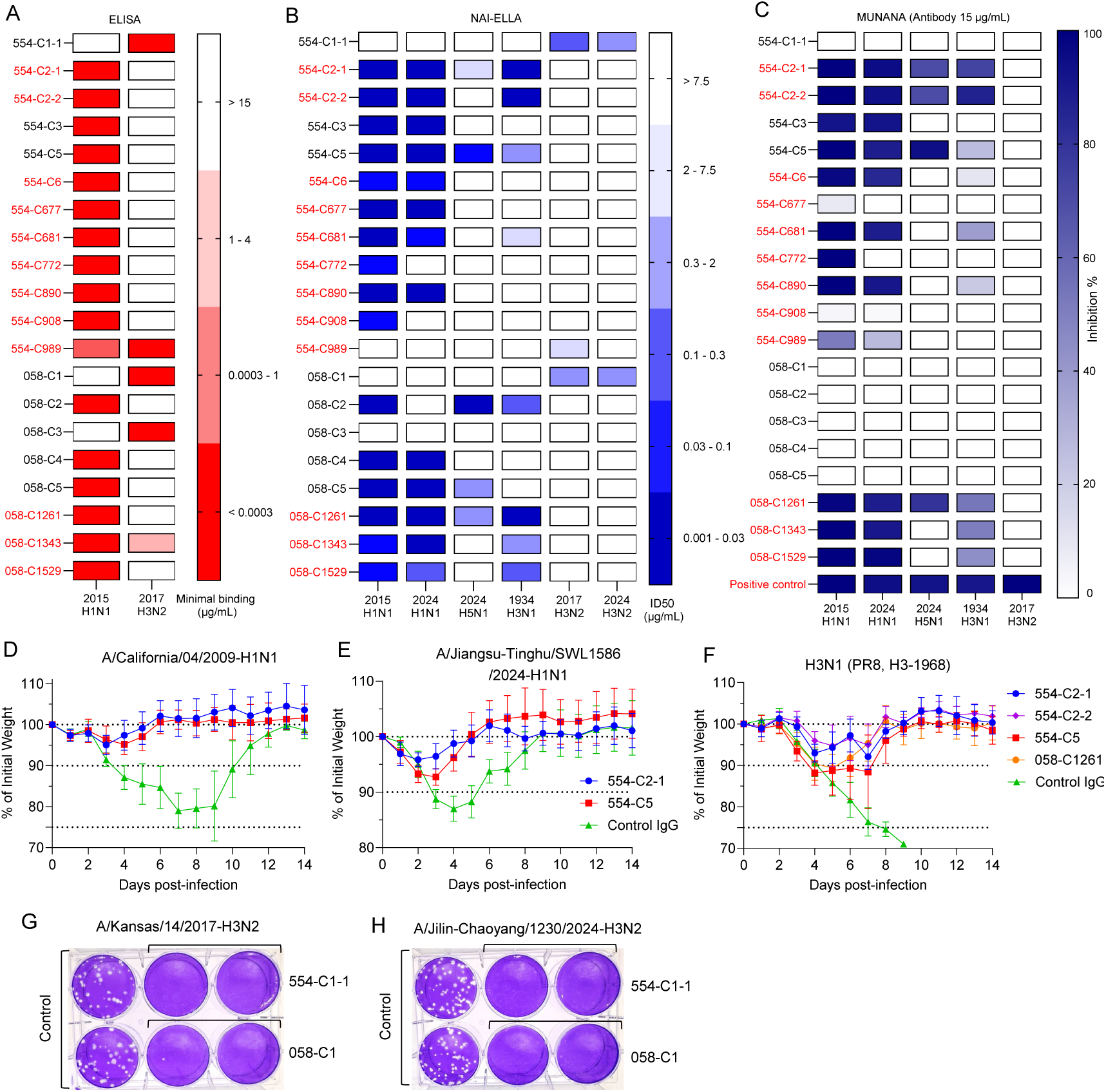
Evaluating the *in vivo* and *in vitro* functionality of NA mAbs isolated from NA-specific MBC repertoires. **(A)** Heatmap showing mAb binding to recombinant N1 (rN1, A/Michigan/45/2015-H1N1) and rN2 (rN2, A/Kansas/14/2017-H3N2) as determined by ELISA. Colored boxes indicate the minimum concentration for detectable binding. **(B)** Heatmap showing mAb NAI titers, as determined by ELLA. Colored boxes represent half–maximal inhibitory dose (ID₅₀). Recombinant NA proteins were used in the ELLA, except that the H3N1 (PR8, H3-1968) virus was utilized. The color boxes indicate the mean of two technical replicates (A and B). (**C)** Heatmap showing mAb inhibition activity (at 15 μg/mL) against H1N1-N1, H5N1-N1, and 2017-H3N2-N2 VLPs, as measured by the MUNANA assay. The H3N1 (PR8, H3-1968) virus was utilized. Colored boxes indicate the mean percentage of inhibition from three technical replicates. **(D to F)** The percentage of initial body weight in BALB/c mice following prophylactic administration of the indicated NA mAbs. Mice (n = 6 per group) received an intraperitoneal injection of the indicated NA mAbs or an irrelevant human IgG (control IgG) 12 hours before being challenged with ∼5 × mLD_50_ of A/California/04/2009 (H1N1) (D) or 1 × 10^6^ PFU of A/Jiangsu–Tinghu/SWL1586/2024 (H1N1) (e), or ∼3 × mLD_50_ of H3N1 (PR8, H3-1968) (F), and the body weight were monitored during the flowing 14 days. The data are presented as mean ± s. d. **(G and H)** MDCK-SIAT1 cells were utilized for plaque reduction assay to measure the inhibitory activity of the indicated NA mAbs against A/Kansas/14/2017(H3N2) and A/Jilin-Chaoyang/1230/2024(H3N2) at 15 μg/mL in the overlay medium.

We evaluated the binding specificity of these mAbs to N1-A/Michigan/45/2015 and N2-A/Kansas/14/2017-H3N2 subtypes and found that 554-C989 and 058-C1343 exhibited cross-reactive binding (Figure 6A, S7A and S7B). Clones 554-C1-1, 058-C1, and 058-C3, derived from the most prevalent clonotypes, bound specifically to N2, whereas the remaining mAbs were specific to N1. Inhibiting activity was assessed using an enzyme-linked lectin assay (ELLA), which revealed that 554-C1-1 and 058-C1 potently inhibited N2 from A/Kansas/14/2017-H3N2 and A/Jilin-Chaoyang/1230/2024-H3N2. In contrast, 554-C989 showed only modest inhibition against the A/Kansas/14/2017-H3N2 N2 strain (Figures 6B, S7C-S7H). Most N1-targeting mAbs demonstrated cross-inhibitory activity against H1N1 strains A/Michigan/45/2015 and A/Jiangsu-Tinghu/SWL1586/2024, except for 554-C772 and 554-C908, which inhibited only A/Michigan/45/2015-H1N1. Notably, 554-C2-1, 554-C5, 058-C2, 058-C5, and 058-C1261 also inhibited N1 from A/dairy-cattle/Texas/24-009495-011/2024-H5N1. Moreover, 554-C2-1, C2-2, C5, C681 and 058-C2, C1261, C1343, C1529 displayed inhibition against historic N1 (PR8-1934), indicating the conservation of N1 epitopes. To determine whether these mAbs targeted the enzymatic site of NA, we performed inhibition assays using 4-(methylumbelliferyl)-N-acetylneuraminic acid (MUNANA). The results showed that most mAbs containing the “DR” motif (red indicated) inhibited neuraminidase activity of two H1N1 strains, with four inhibiting cow-H5N1 and nine inhibiting N1-1934 (Figures 6C and S7I-S7M), indicating recognition of conserved epitopes within the NA catalytic site. However, none of the mAbs inhibited the enzymatic activity of N2 from the modern strain (A/Kansas/14/2017-H3N2).

We next evaluated the *in vivo* protective efficacy of 554-C2-1 and 554-C5 against H1N1 infection (A/California/04/2009 and A/Jiangsu-Tinghu/SWL1586/2024) in mice. Compared to the control group, prophylactic treatment of both mAbs conferred significant protection against morbidity and mortality (Figures 6D, 6E, S7N, and S7O). Additionally, the 554-C2-1, 554-C2-2, 554-C5, and 058-C1261 provided protection against H3N1 (PR8, H3-1968) infection under prophylactic treatment (Figures 6F and S7P). Due to the lack of suitable mouse models for recent human H3N2 isolates, we assessed the inhibitory activity of 058-C1 and 554-C1-1 against H3N2 strains from 2017 and 2024 using a plaque reduction assay. Both mAbs markedly reduced the number and size of plaques formed by the two H3N2 strains (Figures 6G and 6H). These findings demonstrate that mAbs derived from NA-specific MBCs exhibit potent cross-inhibitory activity both *in vitro* and *in vivo*.

### Cryo-EM structure of 554-C2-1 with tetrameric N1 and 554-C1-1 with tetrameric N2

Next, we subject the tetrameric N1 bound with 554-C2-1 and the tetrameric N2 bound with 554-C1-1 to cryo-EM analysis (Figures S8 and S9). Both structures were resolved at an overall resolution of 2.0 Å, assessed by the gold standard of Fourier shell correlation (FSC) = 0.143 (Figures 7A, 7D, S8, S9, and Table S2). These high-resolution structures enable accurate modeling of the antibody-antigen binding interface, assignment of glycans, and positioning of water molecules.

**Figure 7.**
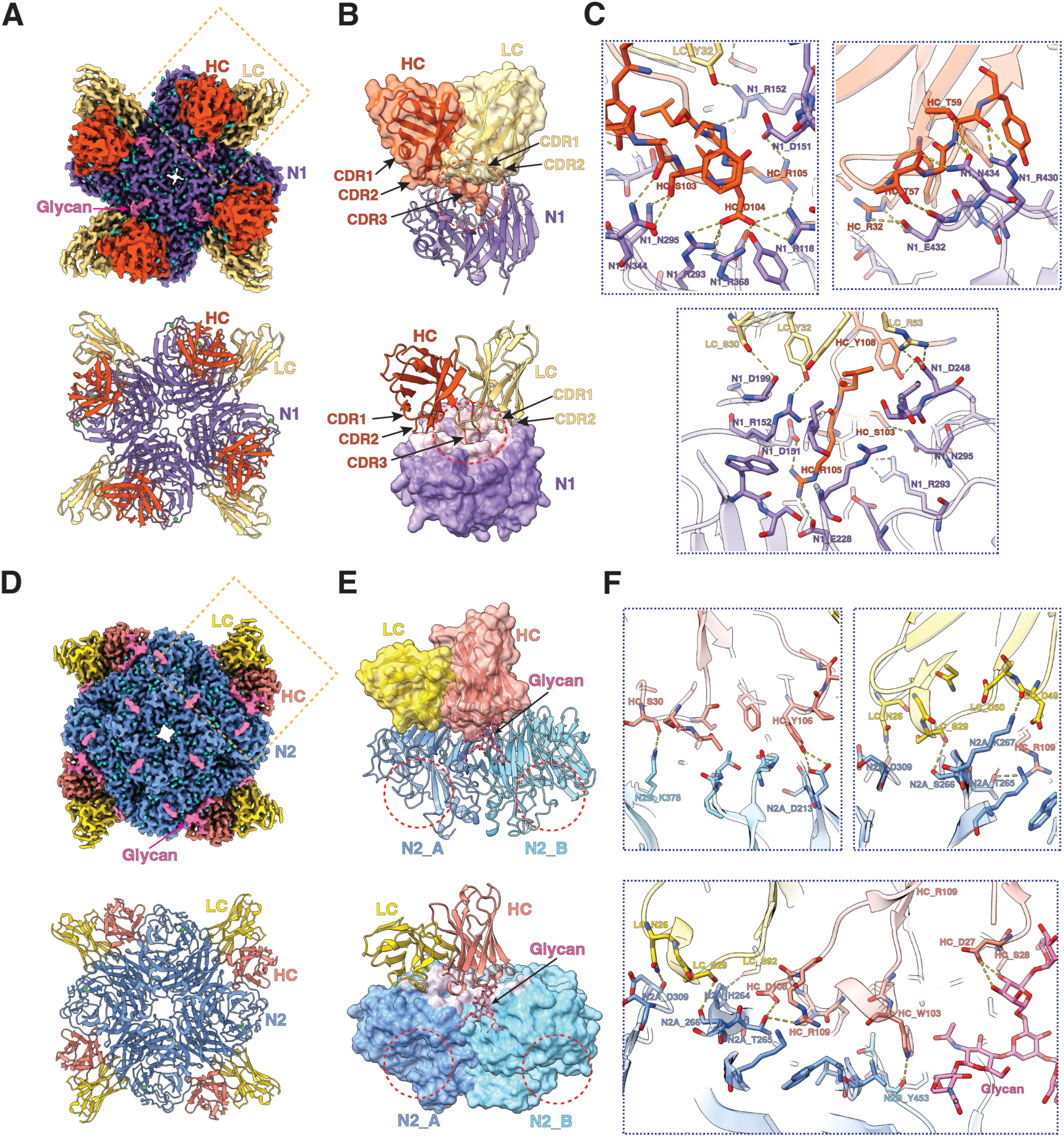
Cryo-EM structures of 554-C2-1 with tetrameric N1 and 554-C1-1 with tetrameric N2. **(A)** The cryo-EM density map and built model of 554-C2-1 with N1 are shown. HC and LC stand for the heavy chain and light chain of the Fab, respectively. The orange box with dashed lines indicates the region for panel (B). **(B)** The upper panel shows the structural model of one Fab interacting with one N1 monomer. HC and LC are shown in cartoon style, overlaid with semi-transparent surface representation. N1 is shown in cartoon style. The circles with red dashed lines mark the N1 enzymatic pocket. The lower panel shows the same view of the model but with N1 shown in surface representation. The N1 residues that interact with the Fab are shown in pink, while the rest are shown in purple. The CDR1/CDR2/CDR3 of HC and the CDR1/CDR2 of LC are indicated by arrows. (**C)** Key interacting residues of HC and LC with N1 are shown as sticks and labeled. Dashed lines indicate hydrogen bonds (H-bonds). **(D)** The cryo-EM density map and built model of 554-C1-1 with N2 are shown. HC and LC stand for the heavy chain and light chain of the Fab, respectively. The orange box with dashed lines indicates the region for panels (E). **(E)** The upper panel shows the structural model of one Fab molecule interacting with two N2 monomers (N2_A and N2_B). HC and LC are shown in cartoon style, overlaid with semi-transparent surface representation. N2 is shown in cartoon style. The circles with red dashed lines mark the N2 enzymatic pocket. The lower panel shows the same view of the model but with N2 shown in surface representation. The N2 residues that interact with the Fab are shown in pink, while the rest are shown in two shades of blue. N200-Glycans are shown as sticks and labeled. **(F)** Key interacting residues of HC and LC with N2 are shown as sticks and labeled. Dashed lines indicate hydrogen bonds (H-bonds).

The cryo-EM structure of the 554-C2-1-N1 complex reveals a previously reported binding mode in which the long HC CDR3 of C2-1 inserts into the N1 active site^35^ (Figure 7B). All CDR loops of C2-1, except CDR3 of LC, contribute to the binding to the active site, effectively blocking the accessibility of the sialic acid substrate. This binding mode and corresponding conserved epitope footprint are consistent with the result of the MUNANA assay. Close examination of the HC CDR3 identifies a “DR” motif (encoded by HC_D104 and HC_R105) (Figure 7C), engaging the critical residues in the active site, such as R118, D151, E228, R293, and R368 of N1. This interaction is further strengthened by an extensive network of hydrogen bonds contributed by other CDRs (Figure 7C).

Interestingly, the structure of the 554-C1-1-N2 complex identifies a previously uncharacterized epitope. This epitope is neither close to the catalytic site nor like the previously described “Dark side” near the stalk region^36,37^. Instead, Fab C1-1 binds to the side of the tetramer, in between the two monomers. A conserved glycan at N200 in the N2 family is also part of this epitope. The elongated epitope, which spans both monomers, engages all CDR loops of C1-1 (Figure 7E). This unique binding mode explains the lack of inhibitory activity in the MUNANA assay for all viral strains tested (Figure 6C). However, antibody binding at the side of the NA tetramer may create steric hindrance, impeding NA activity *in cis* by blocking substrate access on the same membrane. A close-up view of the binding interface reveals fewer hydrogen bonds compared with the 554-C2-1-N1 complex (Figures 7C and 7F). Notably, three residues of HC interact with the glycan, where HC_W103 interacts with glucosamine through electron sharing, and HC_D27/HC_S28 form hydrogen bonds with the mannose residue (Figure 7F).

## Discussion

The NA of the influenza virus, a secondary but relatively conserved glycoprotein, is increasingly recognized as a promising target for next-generation influenza vaccines^10,11,38^. NA-specific immunity in humans has been associated with protection against infection and reduced viral shedding^12,14,16,39^. Here, we developed a membrane-anchored mNA that elicited stronger head-directed and functional antibody responses compared to sNA. In rhesus macaques, mNA induced cross-reactive and cross-protective humoral immunity. Cryo-EM structural analyses revealed conserved epitopes on the N1 and N2 subtypes that are targeted by MBCs generated following mNA immunization. These findings demonstrate that mNA is highly immunogenic and can induce broadly reactive NA-specific immunity, supporting its potential as a foundation for mNA-based influenza vaccines in humans.

Current influenza vaccines elicit limited NA-specific immune responses^40^, likely due to low NA content or improper conformational presentation of NA epitopes^41,42^. We found that mNA exhibited greater head-specific immunogenicity than sNA. The possible explanation is that mNA utilizes its transmembrane domain to form a natural tetramer, avoiding the incorporation of foreign folding domains that could interfere with antigenic head epitopes and impair functional antibody responses. This phenomenon parallels observations in HIV vaccine development, where immunodominant non-neutralizing epitopes in the base region of the envelope glycoprotein (Env) can divert immune responses away from critical neutralizing sites^43,44^. Most seasonal influenza vaccines lack adjuvants, and the commonly used adjuvant Alum fails to enhance human humoral responses to inactivated H5N1 vaccines^45^. Notably, mNA was compatible with both Alum and ISCOM adjuvants, and their inclusion further amplified the humoral response, suggesting broad applicability of the membrane-anchoring strategy across different vaccine platforms.

In humans, NA cross-reactive B cells are characterized by long HCDR3 loops containing a “DR” motif that inserts into the conserved enzymatic pocket of NA^22^. Analysis of the NA-specific BCR repertoires induced by mNA vaccination revealed that the HCDR3 length distribution is distinct from the Gaussian pattern observed in non-antigen-specific B cells in NHPs^46^, with a significant enrichment of BCRs bearing long HCDR3 regions (>17 amino acids). Moreover, many “DR” motif-containing mAbs derived from MBCs demonstrated broad NA inhibitory activity. These findings indicate that mNA vaccination effectively recruits cross-reactive B cells targeting the conserved NA active site. The clone 554-C1, which contains the “DR” motif, exhibited high levels of SHM and diverse clonal evolution, further suggesting that mNA immunization promotes the recruitment of cross-reactive B cell precursors into germinal centers (GCs) for clonal selection and affinity maturation^47,48^. Additionally, mNA vaccination elicited a diverse array of “DR” motif-containing clones, indicating robust engagement of a broad B cell repertoire in the GC response^49^.

Several previous studies have demonstrated that glycosylation at N2, particularly at position N245 in recent H3N2 strains, enables escape from mAbs targeting the enzymatic active site^50–52^. Compared to N1, the magnitude and durability of N2-inhibitory antibody responses were weaker, a phenomenon that may be attributed to increased glycosylation of N2, which obscures functionally important epitopes. Enhancing N2 immunogenicity may therefore require optimizing the N2 immunogen and adjusting the N1:N2 antigen ratio. Despite a decline in circulating N2-inhibitory antibodies over time, we detected N1-specific, N2-specific, and cross-reactive memory B cells (MBCs) two weeks after booster immunization. These MBCs are expected to persist longer than serum antibodies and can be rapidly reactivated upon antigen re-encounter, offering timely protection^53–55^. Notably, we identified that the predominant N2-specific clone elicited by mNA vaccination targets a previously uncharacterized functional epitope located at the interface of two adjacent N2 monomers. This finding indicates that our engineered mN2 preserves tetrameric structural integrity, which is essential for inducing broad N2-inhibitory immune responses and overcoming epitope masking caused by glycosylation of the enzymatic pocket.

## Limitations of the study

Humans exhibit heterogeneous influenza exposure histories. The current study utilized mN1 and mN2 immunization in NHPs to model pre-existing neuraminidase (NA)-specific immunity, although this approach does not fully recapitulate the complexity of human influenza exposure. Although mice receiving serum from mNA-immunized NHPs were protected against influenza infection, we were unable to perform direct challenge experiments in NHPs to assess vaccine-induced protection. Due to the limited availability of NHPs, the immunogenicity of varying mNA doses could not be evaluated. NA broadly reactive MBCs containing the “DR” motif were identified in two immunized NHPs; however, sequencing a larger cohort of NHPs would be necessary to determine the frequency of “DR”-containing NA-specific MBCs and assess their broader relevance.

## Acknowledgements

We thank all the members of the Zeli Zhang laboratory for insightful discussions. We thank the staff at the Laboratory Animal Resources Center and the flow cytometry core at Westlake University for their assistance with techniques. We also appreciate the support from the Cryo-EM facility at Westlake University for Cryo-EM data collection and from the High-Performance Computing Center for computational resources. This project is supported by the Westlake Educational Foundation to Z.Z. and X.Wu, Special Fund of the State Key Laboratory of Gene Expression to (SKLGE-ZX-2025007) X.Wu, Zhejiang Provincial Key Laboratory Construction Project (2024ZY01026) and Platform Development for Novel Vaccines and Antibodies (2024E10060), and Natural Science Foundation of Zhejiang province (LR26H190001) to Z.Z., Zhejiang Key Laboratories Project (2024E10052) to X.Wu, and National Natural Science Foundation of China (NSFC) to Z.Z. (82471855), X.Wang (825B2062), R.S. (82330054), C.L. (82502209), and X.Wu. (32471303).

## Author contributions

Conceptualization: Z.Z., X. Wu; Methodology: R.L., C.L., B.C., Q.Y., X. Wang, and C.C.; Formal analysis: R.L., C.L., B.C., Z.Z., and X. Wu; Investigation: R.L., C.L., and B.C.; Project administration: R.S., Z.Z., and X. Wu; Funding acquisition: Z.Z., X. Wu, and R.S.; Writing: R.L., C.L., Z.Z., and X. Wu; Supervision: Z.Z., X.Wu.

## Supplementary information

Figures. S1 to S9, Tables S1 to S2

## STAR METHODS

### Key resources table

| REAGENT or RESOURCE | SOURCE | IDENTIFIER |
| --- | --- | --- |
| Antibodies |  |  |
| 554-C1-1 | in house | This manuscript |
| 554-C2-1 | in house | This manuscript |
| 554-C2-2 | in house | This manuscript |
| 554-C3 | in house | This manuscript |
| 554-C5 | in house | This manuscript |
| 554-C6 | in house | This manuscript |
| 554-C677 | in house | This manuscript |
| 554-C681 | in house | This manuscript |
| 554-C772 | in house | This manuscript |
| 554-C890 | in house | This manuscript |
| 554-C908 | in house | This manuscript |
| 554-C989 | in house | This manuscript |
| 058-C1 | in house | This manuscript |
| 058-C2 | in house | This manuscript |
| 058-C3 | in house | This manuscript |
| 058-C4 | in house | This manuscript |
| 058-C5 | in house | This manuscript |
| 058-C1261 | in house | This manuscript |
| 058-C1343 | in house | This manuscript |
| 058-C1529 | in house | This manuscript |
| 93(TruStain FcX™) [anti-CD16/32] | Biolegend | Cat#101319<br>RRID: AB_1574973 |
| 2H7 (Brilliant Violet 510) [anti-CD20] | Biolegend | Cat#302339<br>RRID: AB_2561941 |
| SP34-2 (BV650) [anti-CD3] | BD | Cat#563916<br>RRID: AB_2738486 |
| M5E2 (BV650) [anti-CD14] | Biolegend | Cat#301835<br>RRID: AB_11204241 |
| 3G8 (BV650) [anti-CD16] | Biolegend | Cat#302041<br>RRID: AB_11125578 |
| Polyclonal (PE) [anti-IgD] | SouthernBiotech | Cat#2030-09<br>RRID: AB_2795630 |
| G18-145 (BV786) [anti-IgG] | BD | Cat#564230<br>RRID: AB_2738684 |
| RA3-6B2 (PerCP/Cyanine5.5) [anti-CD45R] | Biolegend | Cat#103235<br>RRID: AB_893354 |
| 11-26c.2a (Brilliant Violet 510) [anti-IgD] | Biolegend | Cat#405723<br>RRID: AB_2562742 |
| 281-2 (Brilliant Violet 785) [anti-CD138] | Biolegend | Cat#142534<br>RRID: AB_2814047 |
| SA367H8 (Fas) [anti-CD95] | Biolegend | Cat#152607<br>RRID: AB_2632902 |
| 90 (PE-Cyanine7) [anti-CD38] | Biolegend | Cat#102717<br>RRID: AB_2275531 |
| GK1.5 (Brilliant Violet 711) [anti-CD4] | Biolegend | Cat#100447<br>RRID: AB_2564586 |
| 53-6.7 (FTIC) [anti-CD8a] | Biolegend | Cat#100705<br>RRID: AB_312744 |
| 29F.1A12 (PD-1) [anti-CD279] | Biolegend | Cat#135239<br>RRID: AB_2629767 |
| IM7 (Alexa Fluor700) [anti-CD44] | Biolegend | Cat#103025<br>RRID: AB_2564586 |
| L138D7 (dazzle594) [anti-CD185] | Biolegend | Cat# 145521<br>RRID: AB_2563644 |
| HRP-conjugated goat anti-mouse IgG (H+L) | Invitrogen | Cat#31430<br>RRID: AB_228307 |
| Anti-Monkey IgG (whole molecule)-Peroxidase antibody produced in Rabbit | Sigma-Aldrich | Cat#A2054<br>RRID: AB_257967 |
| HRP-conjugated goat anti-human IgG (H+L) | Jackson ImmunoResearch | Cat#109-035-003<br>RRID: AB_2337577 |
| Bacterial and virus strains |  |  |
| DH5α | in house | N/A |
| DH10bac | in house | N/A |
| A/California/07/2009 (H1N1) | Lab stock | N/A |
| A/Kansas/14/2017 (H3N2) | Lab stock | N/A |
| H3N1 (PR8, H3-1968) | Lab stock | N/A |
| A/Jiangsu-Tinghu/SWL1586/2024 (H1N1) | Lab stock | N/A |
| A/Jilin-Chaoyang/1230/2024 (H3N2) | Lab stock | N/A |
| Biological samples |  |  |
| NHP blood samples | Center for New Drug Safety Evaluation and Research, China Pharmaceutical University | N/A |
| Chemicals, peptides, and recombinant proteins |  |  |
| A/Michigan/45/2015(H1N1) NA | in house | N/A |
| A/Kansas/14/2017(H3N2) NA | in house | N/A |
| A/cattle/Texas/24/2024(H5N1) NA | in house | N/A |
| A/Jilin-Chaoyang/1230/2024(H3N2) NA | in house | N/A |
| A/Kansas/14/2017(H3N2) NA R292K | in house | N/A |
| A/Michigan/45/2015(H1N1) NA R293K | in house | N/A |
| A/Michigan/45/2015(H1N1) NA VLP | in house | N/A |
| A/Kansas/14/2017(H3N2) NA VLP | in house | N/A |
| A/cattle/Texas/24/2024(H5N1) NA VLP | in house | N/A |
| ISCOM | in house | N/A |
| Alhydrogel® adjuvant 2% | InvivoGen | Cat#vac-alu-10 |
| Human FC block | BD | Cat#564200 |
| L/D blue | Thermo | Cat#L34961 |
| Strepavidin-AF647 (AF647) | Biolegend | Cat#405237 |
| Strepavidin-BV421(BV421) | Biolegend | Cat#405226 |
| 7-AAD Viability Staining Solution | Biolegend | Cat#420403 |
| Fetuin A/ASHG human | Sigma-Aldrich | Cat#SRP6217 |
| Bovine serum albumin | BBi | Cat#A600332 |
| Tween-20 | Beyotime | Cat#ST1727 |
| Tween-80 | aladdin | Cat#9005-65-6 |
| CaCl<sub>2</sub> | aladdin | Cat#C110766 |
| HRP-conjugated peanut agglutinin | Sigma-Aldrich | Cat#L7759 |
| PBS | Beyotime | Cat#C0221A |
| TMB substrate | Thermo Fisher | Cat#34021 |
| Dulbecco's Modified Eagle Medium (DMEM) | Gibco | Cat#C11995500BT |
| Roswell Park Memorial Institute (RPMI) 1640 | Gibco | Cat#C11875500BT |
| L-glutamine | Gibco | Cat#A2916801 |
| Fetal bovine serum (FBS) | CellMax | Cat#SA112.02 |
| Penicillin-streptomycin solution | CellMax | Cat#CPS101.02 |
| G418 | Gibco | Cat#10131035 |
| MEM | GENOM | Cat#GNM41502-2 |
| SIM SF | Sino Biological | Cat#MSF1 |
| SIM HF | Sino Biological | Cat#MHF1 |
| DDM | LABLEAD | CAS#69227-93-6 |
| Streptavidin NUPharose FF | Newlong Biology | CAS#NRPB41 |
| Lipofectamine 3000 | Thermo Fisher | Cat#L3000075 |
| PEI transfection reagent | Yeasen, biotechnology | Cat#40816ES03 |
| Uranyl acetate | ELECTRON<br>MICROSCOPY<br>SCIENCES | Cat#22400 |
| Dimethyl sulfoxide (DMSO) | Biosharp | Cat#BS087 |
| His-Tag Protein Purification Resin | Beyotime | Cat#P2218 |
| Protein A Resin | GenScript | Cat#L00210-50 |
| Milk | Beyotime | Cat#P0216 |
| Tris-base | biosharp | Cat#BS083 |
| NaCl | biosharp | Cat#BS112 |
| HCl | Sinopharm | Cat#10011018 |
| Ficoll paque plus | Cytiva | Cat#17144003-1 |
| Glycerol | Sigma-Aldrich | Cat#56-81-5 |
| Agarose | BBI | Cat#A600015-0005 |
| diethylaminoethyl (DEAE) | Yuanye | Cat#S14097 |
| L-1-tosylamido-2-phenylethyl chloromethyl ketone (TPCK) | Worthington Biochem | Cat#LS003740 |
| paraformaldehyde (PFA) | Biosharp | Cat#BL539A |
| crystal violet | Beyotime | Cat#C0121 |
| Biotin Labeling Kit for Avi-tag Protein with BirA | Beyotime | Cat#P0630S |
| ELISA Stopping Solution | BBI | Cat#E661006 |
| Critical commercial assays |  |  |
| NA-Fluor™ Influenza Neuraminidase Assay Kit | Applied Biosystems | Cat#4374422 |
| Deposited data |  |  |
| Cryo-EM map of the 554-C2-1-N1 complex | Electron Microscopy<br>Data Bank<br>https://www.ebi.ac.uk/emdb/ | EMD-67987 |
| Structural model of the 554-C2-1-N1 complex | Protein Data Bank<br>https://www.rcsb.org/ | 21TM |
| Cryo-EM map of the 554-C1-1-N2 complex. | Electron Microscopy<br>Data Bank<br>https://www.ebi.ac.uk/emdb/ | EMD-67988 |
| Structural model of the 554-C1-1-N2 complex. | Protein Data Bank<br>https://www.rcsb.org/ | 21TN |
| Experimental models: Cell lines |  |  |
| Hi5 cells | Sino Biological | N/A |
| Sf9 cells | Sino Biological | N/A |
| MDCK-SIAT1 | Sigma-Aldrich | N/A |
| MDCK cells | ATCC | N/A |
| HEK293F cells | ATCC | N/A |
| Experimental models: Organisms/strains |  |  |
| Mice: BALB/c | Westlake University<br>Vivarium | N/A |
| NHP: Rhesus macaques | Center for New Drug<br>Safety Evaluation and<br>Research, China<br>Pharmaceutical<br>University | N/A |
| Recombinant DNA |  |  |
| pMDLg/pRRE | Addgene | Cat#12251 |
| pRSV-Rev | Addgene | Cat#12253 |
| pcDNA3.1-NA | This manuscript | N/A |
| pFastBac-NA | This manuscript | N/A |
| Software and algorithms |  |  |
| FlowJo 10.8.1 | FlowJo, LLC | www.flowjo.com |
| GraphPad Prism 10 | GraphPad | www.graphpad.com |
| Excel | Microsoft | N/A |
| cryoSPARC v4 | Structura<br>Biotechnology Inc. | https://cryosparc.co<br>m/ |
| UCSF Chimera |  | https://www.cgl.ucsf.<br>edu/chimera/ |
| AlphaFold3 |  | https://alphafoldserv<br>er.com/ |
| Coot |  | https://www.ccp4.ac.<br>uk/ |
| Phenix |  | https://phenix-online.org |
| UCSF ChimeraX |  | https://www.cgl.ucsf.<br>edu/chimerax/ |
| Pymol | Schrodinger. | https://pymol.org/2/ |
| 10X Genomics CellRanger pipeline V7.1.0 |  | https://www.10xgeno<br>mics.com/ |
| MEGA V11 |  | https://www.megasof<br>tware.net/ |
| iTOL |  | https://itol.embl.de/ |
| IgBLAST |  | https://www.ncbi.nlm.nih.gov/projects/igblast/ |
| IMGT |  | https://www.imgt.org/ |
| Other |  |  |
| 96-well black plates | Thermo | Cat#237105 |
| Flat bottom 96-well plates | Thermo | Cat#168055 |
| 96-half-well plates | Corning | Cat#3690 |

## Resource availability

### Lead contact

Further information and requests for resources and reagents should be directed to and will be fulfilled by the lead contact, Zeli Zhang.

### Materials availability

Reagents generated in this study include NHP anti-NA mAbs and recombinant NA proteins. DNA expression constructs for the mAbs and recombinant NA proteins are available upon request.

### Data and code availability

The cryo-EM density maps and atomic coordinates have been deposited in the Electron Microscopy Data Bank (EMDB) and Protein Data Bank (PDB) under accession numbers EMD-67987 and PDB: 21TM, for the 554-C2-1-N1 complex; EMD-67988 and PDB: 21TN, for the 554-C1-1-N2 complex.

### Experimental model and study participant details Mice and NHPs

Mouse experiments were conducted at Westlake University. All experimental procedures were approved by the Laboratory Animal Resources Center (LARC) at Westlake University (Approval No. AP#23-037-ZZL-10). Experiments were performed using sex- and age-matched female mice aged 7 to 8 weeks. BALB/c mice (strain 000651) were obtained from Shanghai Lingchang Biotechnology Company. Rhesus macaques (Macaca mulatta) were housed, immunized, and sampled at the Center for New Drug Safety Evaluation and Research, China Pharmaceutical University. All procedures involving non-human primates were performed in accordance with the Guidelines for the Husbandry and Management of Laboratory Animals at China Pharmaceutical University and were approved by the Institutional Animal Care and Use Committee of China Pharmaceutical University. Eight rhesus macaques (three males and five females) that met quarantine and acclimatization requirements were included in the study. Eight rhesus macaques were screened that had no influenza NA background immunity, as determined by ELISA. Ethical approval for this study was granted under license number I2024015.

### Cell lines

HEK293T and MDCK cells were cultured in Dulbecco’s Modified Eagle Medium (DMEM) (Gibco) supplemented with 10% fetal bovine serum (FBS) (CellMax), 2 mM L-glutamine (Gibco), and 100 U/ml penicillin-streptomycin solution (CellMax). Expi293 cells were cultured in a serum-free Chemically Defined Medium (Union Bio). Sf9 cells were cultured in serum-free insect SF-9 Cell Medium (SIM-SF, Sino Biological) supplemented with 100 U/ml penicillin-streptomycin. Hi5 cells were cultured in serum-free insect Hi-5 Cell Medium (SIM-HF, Sino Biological) supplemented with 100 U/ml penicillin-streptomycin. MDCK-SIAT1 cells (Sigma-Aldrich) were maintained in MEM supplemented with 10% FBS, 2 mM L-Glutamine, 100 U/ml of Penicillin-Streptomycin, and 1 mg/ml G418.

### Influenza virus

The viruses A/California/04/2009 (H1N1) and A/Kansas/14/2017 (H3N2) were obtained from our laboratory stock. H3N1 (PR8, H3-1968) was resorted with PR8 backbone expect H3 was derived from A/Aichi/2/1968. A/Jiangsu-Tinghu/SWL1586/2024 (H1N1) and A/Jilin-Chaoyang/1230/2024 (H3N2) were provided by the National Influenza Center at the Chinese Center for Disease Control and Prevention (China CDC). A/Kansas/14/2017 (H3N2) and A/Jilin-Chaoyang/1230/2024 (H3N2) were propagated in MDCK-SIAT1 cells at 35 °C for 48 hours, while the remaining viruses were propagated in MDCK cells at 37 °C for 48 hours. Viral supernatants were clarified by centrifugation, aliquoted, and stored at −80 °C. Viral stock titers were determined using a standard plaque assay.

## Method details

### Expression and purification of sNA and mNA

The sN1 (A/Michigan/45/2015, H1N1) and sN2 (A/Kansas/14/2017, H3N2) with tetrabrachion (TB) or VASP tetramerization domain were expressed and purified as previously described^20^. To obtain mNA immunogens, the full-length N1 or N2 was cloned into pFastBac with an N-terminal Strep-tag. The mN1 and mN2 were expressed in Sf9 cells via recombinant baculoviruses. Specifically, the P0 of recombinant baculoviruses was passaged in Sf9 cells twice to obtain a high viral titer (P2 stock). Then, the Sf9 cells were infected with MOI=5 of P2 viral stock, and the mNA-expressing Sf9 cells were harvested 3 days post-infection (50-60% cell viability) and lysed in a buffer of 25 mM Tris-HCl (pH 7.5), 150 mM NaCl, 2mM CaCl_2_, and 1% DDM. The lysate was subjected to affinity purification using Streptavidin NUPharose FF (Newlong Biology) beads, and further purification was performed by size-exclusion chromatography (SEC) on a Superose 6 or Superdex 200 (10/300 GL) (Cytiva) column equilibrated with 25 mM Tris-HCl (pH 7.5), 150 mM NaCl, 2mM CaCl_2_, 5% glycerol, and 0.03% DDM. Fractions corresponding to the tetrameric form of NA were collected. The purified mNA protein was subjected to several freeze-thaw cycles, and its structural integrity after freeze-thaw was assessed via SEC and SDS-PAGE analysis.

### Negative stain electron microscopy

The sNA protein was diluted to a concentration of 0.004–0.02 mg/mL in 25 mM Tris (pH 7.5), 150 mM NaCl, and 2 mM CaCl₂. The mNA protein was similarly diluted to 0.004–0.02 mg/mL in the same buffer supplemented with 0.02% PS80. Samples were adsorbed onto glow-discharged carbon-coated copper grids and stained with 1% uranyl acetate. Data were acquired using a 120 kV transmission electron microscope (CRYO-EM005).

### Animal immunization design

Seven to eight-week-old female BALB/c mice were used to evaluate the immunogenicity of mNA and sNA. Mice were vaccinated with indicated doses of mNA or sNA protein, formulated with or without Alum or ISCOM adjuvants. PBS was the formulation buffer for sNA, and PBS with 0.02% PS80 was the formulation buffer for mNA. Blood samples were collected before immunization and 14 days after the second immunization. For non-human primates’ immunization, eight rhesus macaques (Macaca mulatta) were randomly assigned to two groups (one male and two females for Alum group-G1 and two males and three females for ISCOM group-G2) to evaluate the immunogenicity of mNA. G1 received 30 μg mNA with 300 μg Alum per animal. G2 received 15 μg mNA with 50 μg ISCOM per animal. To model NA pre-immunization, animals were initially immunized with either N1 or N2 alone, followed by a combined immunization. In the ISCOM group, the adjuvant amount remained at 50 μg per animal, whereas in the Alum group, it was increased to 600 μg per animal for the combined immunization. Immunizations were administered intramuscularly at a single site (the left thigh) with a total volume of 0.5 mL per injection. Blood was collected before vaccination and on days 21, 42, 63, 77, 91, 109, and 330. Plasma was separated, and PBMCs were isolated for downstream analyses.

### Influenza NA ELISA

Mouse serum: High-binding 96-well half-area plates (Corning #3690) were coated overnight at 4°C with recombinant NA protein (VASP-NA) diluted to 1 μg/mL in PBS. Plates were blocked for 1 h at room temperature with PBS containing 0.1% Tween-20 (PBST) and 3% bovine serum albumin (BSA; BBI Life Sciences). After six washes with TBST, plates were incubated with HRP-conjugated goat anti-mouse IgG (H+L) (Invitrogen; 1:10,000) for 90 min at RT. Following final washes, 50 μL per well of TMB substrate was added for 5 min at RT, and reactions were stopped with 50 μL per well stop solution (BBI Life Sciences). Absorbance was read at 450 nm using a microplate reader. Data were analyzed using Excel and GraphPad Prism, with positivity defined as an optical density (OD) at 450 nm of≥ 0.1. NHP plasma and mAbs: Plates were coated with recombinant NA protein diluted to 2 μg/mL. Plates were blocked with PBST containing 3% skim milk (Beyotime) and 2% BSA. Monoclonal antibody detection plates were alternatively blocked with 3% BSA in PBST under identical conditions. After six TBST washes, plates were incubated with the appropriate HRP-conjugated secondary antibody: rabbit anti-monkey IgG (whole molecule)-HRP (Sigma; 1:40,000) for NHP plasma, or HRP-conjugated goat anti-human IgG (H+L) (Jackson ImmunoResearch; 1:10,000) for mAbs, each for 90 min at RT. After final washes, 50 μL per well of TMB substrate was added for 5 min at RT, and reactions were stopped with 50 μL per well stop solution. Absorbance was measured at 450 nm, and data were analyzed as above with the same positivity threshold.

### Enzyme-linked Lectin Assay (ELLA)

MaxiSorp 96-well plates (Thermo Scientific) were coated with fetuin (50 μg/mL in PBS, 150 μL/well, Sigma) overnight at 4°C. After blocking with 1% BSA (BBI Life Sciences) in PBS (200 μL/well, 1 h, RT), plates were washed with TBST (0.1% Tween-20) and incubated with serially diluted recombinant NA or influenza virus (3-fold dilutions in PBS with 1 mM CaCl2 and 0.1% BSA) for 16 h at 37°C. Post-wash, 100 μL/well of HRP-conjugated peanut agglutinin (2 μg/mL, Sigma) was added (1.75 h, RT, dark). Following TBST washes, TMB substrate (100 μL/well) was reacted for 5 min before termination with stop solution (BBI Life Sciences). Absorbance (450 nm) was measured, and NA/virus quantities yielding 90-95% maximal signal were selected for inhibition assays. For NA inhibition (NAI) testing, antibodies or heat-inactivated plasma were serially diluted (starting at 30 μg/mL or 1:10) in PBS containing CaCl_2_ and pre-incubated with standardized NA/virus. Mixtures were transferred to fetuin-coated plates and processed as above. Positive controls (virus/NA without inhibitors) occupied eight wells per plate. ID50 values were calculated as the antibody/serum concentration that achieved a 50% reduction in NA activity relative to controls (GraphPad Prism 10). Samples failing 50% inhibition at the lowest dilution (1:10) were assigned a titer of 10.

### Detection of NA-specific GC B and plasmablast cells in mice

Female BALB/c mice (n = 5 per group) were subcutaneously immunized with 62.5 µg of Alum adjuvant combined with either 2.5 μg of mNA or sNA. Eight days post-immunization, draining inguinal lymph nodes (iLN) were isolated for flow cytometry analyses. Single-cell suspensions were prepared by mechanical dissociation. Antibody staining was performed in FACS buffer. Cells were first pre-incubated with TruStain FcX™ (anti-mouse CD16/32) antibody for 20 min at 4 °C, followed by stained with 300 ng of biotinylated mutant N2 (R292K) with fluorophore-conjugated streptavidin-AF647 or streptavidin-BV421 probes at 4 °C for 30 mins. Without washing, cells were then stained with a surface antibody cocktail (CD19, CD4, CD44, CD8a, IgD, CD138, Fas, CD38, CXCR5, PD-1,) at 4 °C for 30 mins. Dead cells were labeled using Live/Dead Blue. Data acquisition was performed using Cytek Aurora 3 and analyzed using FlowJo 10.

### Flow cytometry and sorting

To detect NA-specific memory B cells in NHP PBMC, frozen PBMC were thawed and recovered in RPMI supplemented with 10% FBS. The recovered cells were counted and stained with the appropriate staining panel. NA (N1 and N2)-specific memory B cells (MBCs) were identified using fluorochrome-labeled N1 (R293K) and N2 (R292K) mutant probes. Fluorescent antigen probes were prepared by mixing 300 ng of biotinylated mutant NA with fluorophore-conjugated streptavidin-Alexa Fluor 647 or streptavidin-BV421 for 30 min at 4°C. For each sample, 1-5 million cells suspended in 50 µL of FACS buffer (0.5%BSA in PBS) were pre-incubated with 2 µg Human BD Fc Block™ for 20 min at 4 °C, followed by staining with NA probes for 4 °C for 30 mins. Without washing, cells were subsequently incubated with a surface antibody cocktail (CD3, CD14, CD16, CD20, IgD, IgG) for 30 min at 4 °C. After washing, cells were resuspended in FACS buffer containing diluted 7-AAD viability dye and acquired on a BD FACSAria Fusion cytometer. The downstream data analysis was performed using FlowJo software version 10.8.2. The frequency of NA-specific MBCs was calculated as a percentage of total MBCs (lymphocytes, singlets, live, CD3⁻CD14⁻CD16⁻CD20⁺IgD⁻IgG⁺^/^⁻) according to the gating strategy shown in Fig. S4. To perform single NA-specific BCR sequencing, 15,000 and 8,000 NA-specific memory B cells were sorted in NHP-058 and NHP-554, respectively, using the above staining and gating strategy.

### BCR sequencing and analysis

Sorted NA-specific MBCs were loaded directly onto a 10X Chromium A chip. Single-cell lysis and cDNA synthesis were performed using the 5’ library and gel bead kit per the manufacturer’s protocol. Custom primers targeting NHP immunoglobulin constant regions were included during V(D)J amplification. Libraries were sequenced on an Illumina NovaSeq platform following standard procedures. Raw FASTQ files were processed using the 10X Genomics CellRanger pipeline (version 7.1.0). Assembled contigs were annotated using IgBLAST and IMGT with a custom rhesus macaque reference.

### Construction of the NA mAbs clonal evolutionary tree

Clonal evolutionary trees were generated as described^22^. Briefly, germline genes were reconstructed by replacing the CDR3 loops in germline sequences with those from B cell clones exhibiting the lowest VH mutation rate within each clonotype, as the original junctional sequences could not be unambiguously inferred. Heavy-chain VDJ sequences were aligned, and phylogenetic trees were constructed in MEGA (v11) with the neighbor-joining method. Trees were rooted to the closest germline immunoglobulin genes and then annotated and visualized in iTOL.

### Generation of NA-VLP

NA-containing virus-like particles (NA-VLPs) were produced by co-transfecting HEK293T cells with 750 ng of pMDLg/pRRE, 250 ng of pRSV-Rev, and 800 ng of pcDNA3.1 plasmid encoding the indicated NA using Lipofectamine 3000 (Thermo) following the manufacturer’s protocol. Cell culture supernatants were collected 48 h post-transfection and stored at −80°C in aliquots until use.

### NA-MUNANA assay

The NA-inhibition activities of mAbs were measured by the NA-Fluor™ Influenza Neuraminidase Assay Kit. Briefly, monoclonal antibodies (15 µg/ml) were incubated with a constant amount of either NA-VLP or influenza virus in 96-well black plates (Thermo) at 37 °C for 1 h. The reaction was then initiated by adding the NA-Fluor™ substrate (MUNANA) to a final concentration of 200 μM and allowed to proceed for an additional hour at 37 °C. Enzymatic activity was terminated with NA-Fluor™ stop solution, and fluorescence signals were recorded on a Synergy H1 microplate reader (BioTek) with excitation at 365 nm and emission at 445 nm.

### *In vivo* challenge experiments

To evaluate whether sera from immunized NHPs confer protection in mice at different time points, sera from the Alum and ISCOM groups at matching time points (D0, D77, D109) were pooled by combining 500 μL from each animal within the group. The pooled sera were transferred (400 µL per mouse) to mice via intraperitoneal injection to determine inhibitory activity against A/California/04/2009-H1N1 (5×mLD50 per mouse), A/Jiangsu-Tinghu/SWL1586/2024-H1N1 (1×10^6 PFU per mouse), and H3N1 (PR8, H3-1968) (3×mLD50). Two additional groups that received an equivalent volume of PBS served as negative controls. Additionally, eight- to ten-week-old mice were used to evaluate the prophylactic potential of NA-specific mAbs. All mAbs were diluted in 200 μL PBS and administered intraperitoneally. Negative control groups received an irrelevant human IgG mAb at equivalent doses. Prophylaxis: mice received 200 μg of the indicated NA mAbs on day 0. Twenty-four hours later (day 1), mice were anesthetized with pentobarbital and intranasally challenged with A/California/04/2009-H1N1 (5×mLD50), A/Jiangsu-Tinghu/SWL1586/2024-H1N1 (1×10^6 PFU), and H3N1 (PR8, H3-1968) (3×mLD50). Body weight was monitored daily for 14 days; mice losing more than 25% of their initial body weight were euthanized in accordance with ethical guidelines.

### The mAb and Fab expression and purification

The Expi293F cell expression system was used for the expression of monoclonal antibodies. The VDJ fragments of the heavy and light chains were cloned into IgG1 and IgK or IgL expression vectors, respectively. The heavy-chain and light-chain expression plasmids were co-transfected into Expi293F cells at a ratio of 1:1 using PEI transfection reagent (Yeasen, biotechnology). The antibodies were purified from the culture supernatant using Protein A agarose resin (GenScript). The purified antibodies were subjected to buffer exchange and concentration and then sterilized by 0.22 μm filtration. For the Fab expression, the heavy-chain Fab fragment of the indicated mAbs was cloned into the pcDNA3.1 vector with a C-terminal His tag, and the expression plasmid was co-transfected with the light-chain expression plasmid into Expi293F cells at a ratio of 1:1 using PEI transfection reagent (Yeasen, biotechnology). The secreted recombinant Fab was purified using His-tag protein purification resin (Beyotime, Biotechnology) and further purified by SEC (Superose 6 or Superdex 200, Cytiva) to remove impurities and multimers, resulting in high-purity Fab fragments. Finally, the Fab fragments were sterilized by 0.22 μm filtration.

### Plaque reduction neutralization assay

MDCK-SIAT1 cells (5 × 10^5^ cells/well) were seeded in 6-well plates and cultured for 24 hours. The indicated influenza viral strains were incubated with mAbs (15μg/mL) for 1 hour at room temperature (RT). Cells were washed with PBS and infected with the virus-mAb mixture for 1 hour at 37°C. A 2 mL overlay mixture was added. Cells were incubated for 48-72 hours at 37°C. Following fixation with 4% paraformaldehyde (PFA) for 20 minutes at RT, the overlay was removed, and plaques were visualized via crystal violet staining (20 minutes at RT). Plaques were counted after washing with sterile water to assess the efficacy of mAb inhibition.

### Cryo-EM Sample Preparation and Data Collection

To form a complex of NA-Fab, purified NA (∼1mg/mL) was incubated with purified Fab (∼1mg/mL) at a molar ratio of 1:1 on ice for 30 mins. The complex was modified with low-molecular-weight polyethylene glycol (PEG) at surface-exposed lysine residues to reduce aggregation and dissociation of the complex on the grids^56^. Specifically, the complex at ∼1 mg/mL was incubated with MS(PEG)12 methyl-PEG-NHS-ester (Thermo Fisher Scientific) at a molar ratio of 1:30 for 2 hrs on ice. Then, 3 μL of the treated complex was applied to glow-discharged (Coolglow, SuPro Instruments) holey carbon grids (Quantifoil R1.2/1.3 400 mesh, Au). The grids were blotted for 4 seconds at ∼100 % humidity and 4 °C, and then plunge-frozen in liquid ethane using the Vitrobot Mark IV (Thermo Fisher Scientific). The cryo-EM data were collected on a Titan Krios G4 (FEI) operating at 300 kV, equipped with a Selectris X imaging filter (Thermo Fisher Scientific) and a Falcon 4i direct electron detector (Thermo Fisher Scientific). An imaging filter with a slit width of 10 eV was used. All cryo-EM movies were recorded using EPU (Thermo Fisher Scientific), with a total electron dose at ∼50 electrons/Å2. The nominal magnification of 130 kx corresponds to a calibrated pixel size of 0.92 Å on the specimen. The defocus range for the samples was between 1.1 and 2.2 µm.

### Image processing

The image processing workflows are illustrated in detail in Fig. S8-9 and were performed using cryoSPARC v4^57^. Briefly, motion-correction and dose weighting were performed using the Patch_Motion_Correction module in cryoSPARC v4. After CTF was estimated with the Patch_CTF module, micrographs with CTF-estimated resolution worse than 4 Å, high drift values, and extreme defocus range were excluded from further data analysis. Data analysis was conducted iteratively. Initial particles from 1000 micrographs were picked using the Blob_Picker module^58^. Two-dimensional (2D) classifications were performed on these picked particles, and 2D averages showing nice protein features were used as initial templates for particle picking on the 1000 micrographs using the Template_Picker module. After 2 rounds of 2D classifications to remove junk particles and false picks, particles belonging to the nice 2D class averages were selected for an Ab-initio 3D reconstruction using C4 symmetry. Clear secondary structure features were readily visible in the initial 3D reconstruction. This initial 3D volume was then utilized to generate 20 2D templates using the Create_Templates module. These generated 2D templates were used for particle picking across all selected micrographs. After 2 rounds of 2D classifications, particles of good quality were subjected to one round of heterogeneous refinement using one good 3D volume and one bad 3D volume, using C4 symmetry. Non-uniform refinement was performed on the resulting good class using C4 symmetry, followed by Local_CTF_refinement and reference-based motion correction. The CTF-refined and polished particles were subjected to another round of non-uniform refinement using C4 symmetry. Local resolution variations were estimated in cryoSPARC. Resolutions of the refinement were calculated according to the gold-standard Fourier Shell Correlation (FSC) 0.143 criterion.

### Model building and refinement

AlphaFold 3 predictions^59^ of the N1, N2, and corresponding Fabs were used as initial models for rebuilding the model into the cryo-EM density map at high resolution. The initial models were docked into the cryo-EM density map using UCSF ChimeraX^60^. Iterative manual rebuilding and adjustment were performed in Coot^61^, including additions of water molecules, glycans, and ions. Then, the model was refined using Phenix^62^. Structural figures were prepared using ChimeraX^63^.

### Quantification and statistical analysis

The statistical analysis methods are indicated in each figure legend.

