## Supplemental Figures for "Membrane-anchored influenza neuraminidase vaccine drives human-like broadly protective B cell responses"

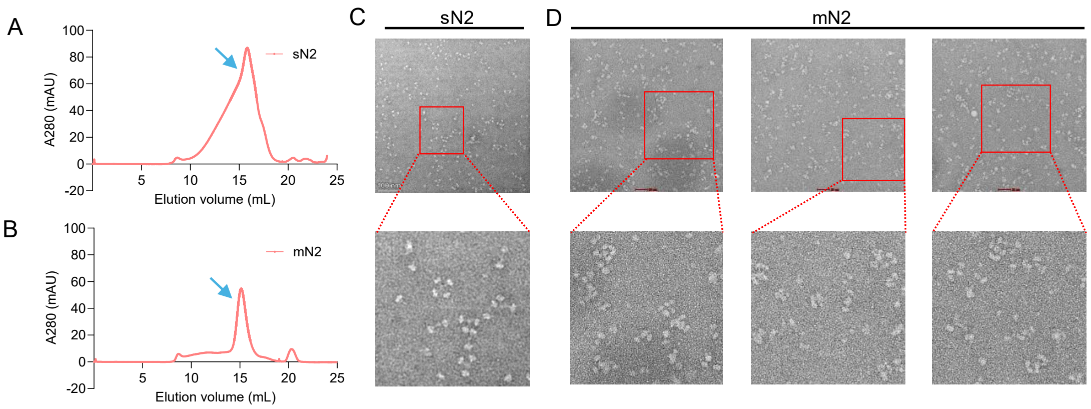


**Figure S1. Expressing the sN2, mN2, and comparing their immunogenicity in mice. (A and B)** Size-exclusion chromatography (SEC) profiles of sN2 (A) and mN2 (B) analyzed on a Superose 6 Increase 10/300 GL column. **(C and D)** Negative staining-EM analysis of purified sN2 (C) and mN2 (D).

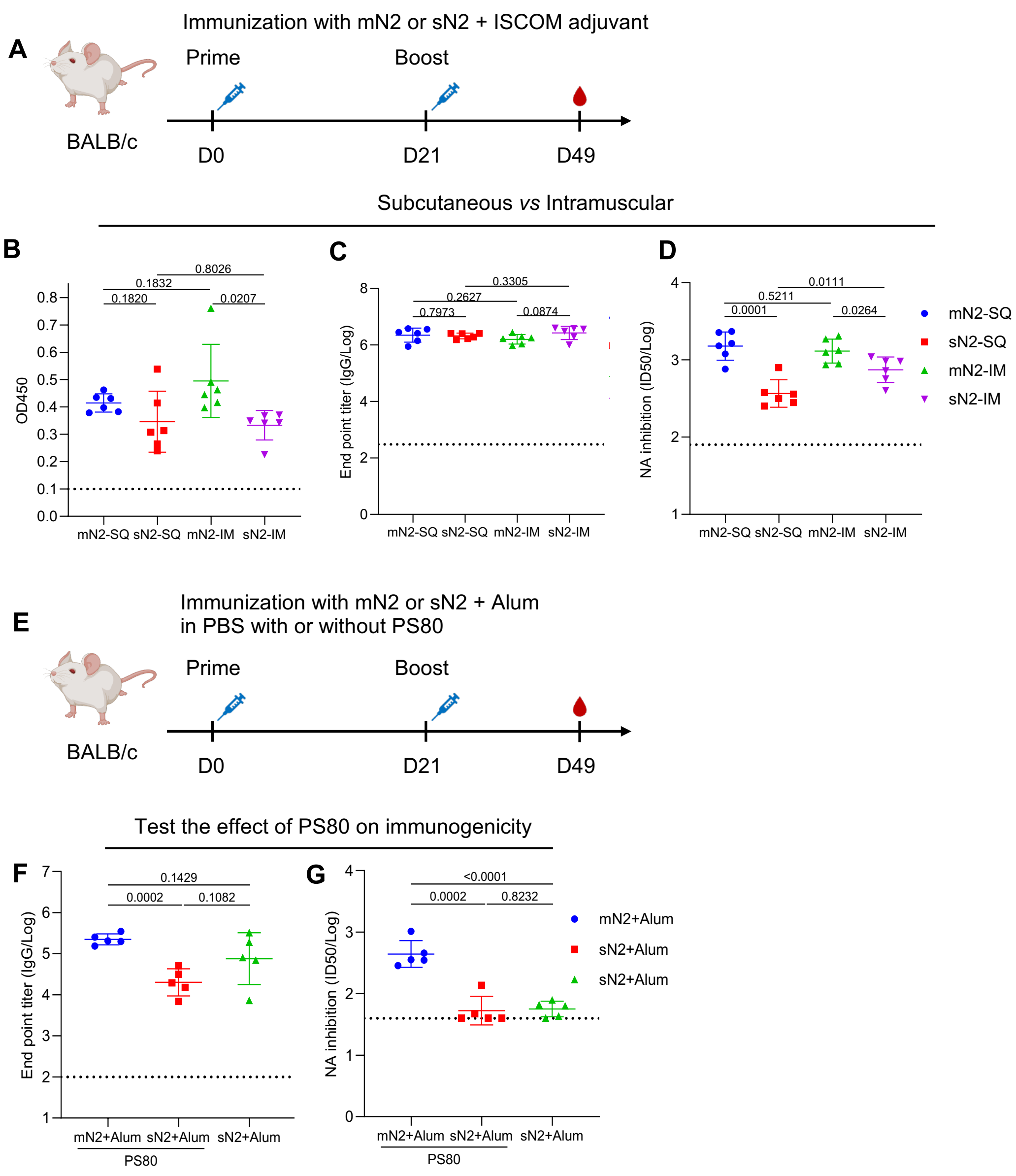


**Figure S2. Comparing the immunogenicity of sN2 and mN2 in mice. (A to D)** BALB/c female mice (n = 6 per group) were immunized via intramuscular or subcutaneous routes at the indicated time with sN2 or mN2. Antibody levels against the whole virus (A/Kansas/14/2017-H3N2) were determined by virus-coated ELISA (B). NA head-specific IgG titers were determined by recombinant VASP-NA-coated ELISA (C). NI titers were determined by ELLA (D). **(E to G)** BALB/c female mice (n = 5 per group) were prime-boosted via subcutaneous routes with mNA or sNA formulated with the PS80. N2 head-specific IgG titers (F) or NAI titers (G) were shown. Data were log-transformed (C, D, F and G) presented as mean ± s. d. Statistical analyses were performed by two-tailed unpaired t-tests, and the corresponding *p*-values were indicated.

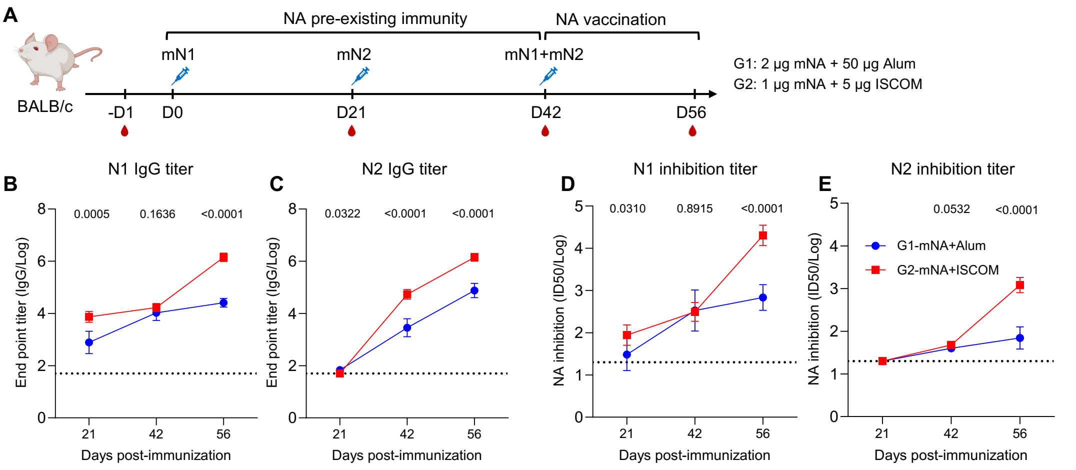


**Figure S3. Establishing NA pre-existing immunity in mice. (A to E)** Female BALB/c mice (n = 6 per group) were immunized with mNA at the indicated dose and time. Group 1 (G1) received 2 μg mNA with Alum adjuvant. Group 2 (G2) received 1 μg mNA with ISCOM adjuvant. NA head-specific IgG antibody levels in sera were measured by ELISA (B and C). NAI titers were determined by ELLA (D and E). Data were log-transformed and presented as mean ± s. d. Statistical analyses were performed on log-transformed data using unpaired *t*-tests (B to E), and the corresponding *p*-values were indicated.

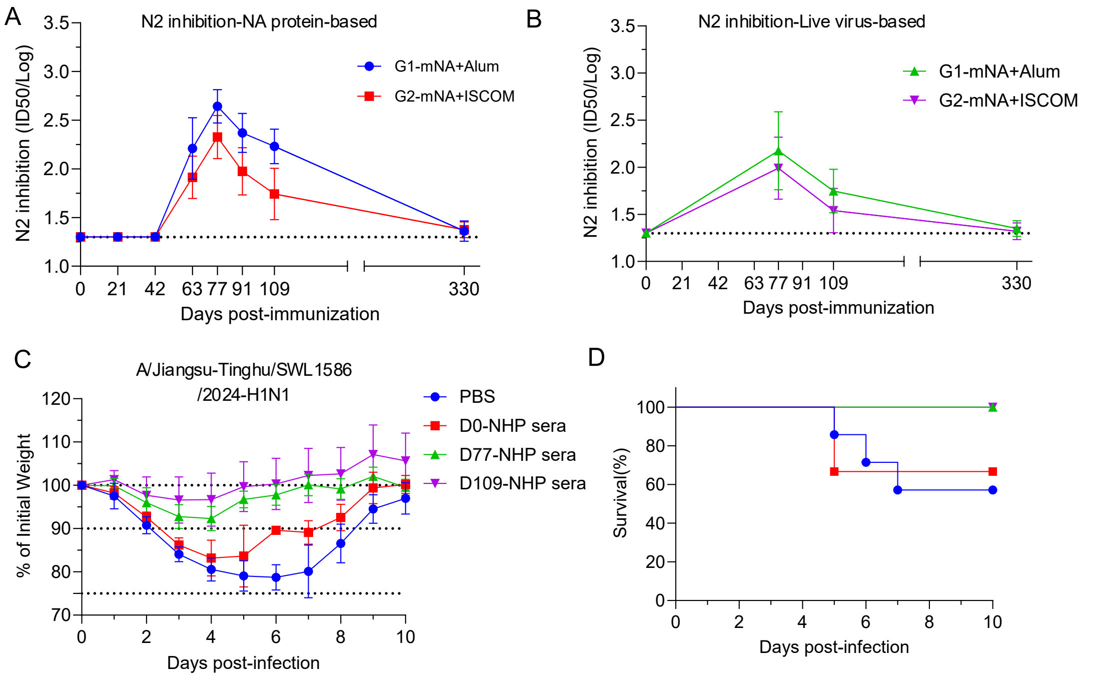


**Figure S4. The immunogenicity and protection of mNA in NHPs. (A and B)**

N2-NAI titers in NHP immune sera were determined by recombinant sN2 (A) or the wild-type virus (A/Kansas/14/2017-H3N2) (B). **(C and D)** The percentage of initial body weight and survival rates in BALB/c mice following passive transfer of the indicated NHP immune sera (eight NHP sera were equally pooled). Mice (n = 6 per group) received an intraperitoneal injection of the indicated NHP immune sera or PBS, 12 hours post sera-transferring, mice were challenged with $1 \times 10^6$ PFU of A/Jiangsu-Tinghu/SWL1586/2024 (H1N1). The percentage of body weight and the survival rate were shown. The data are presented as mean ± s. d.

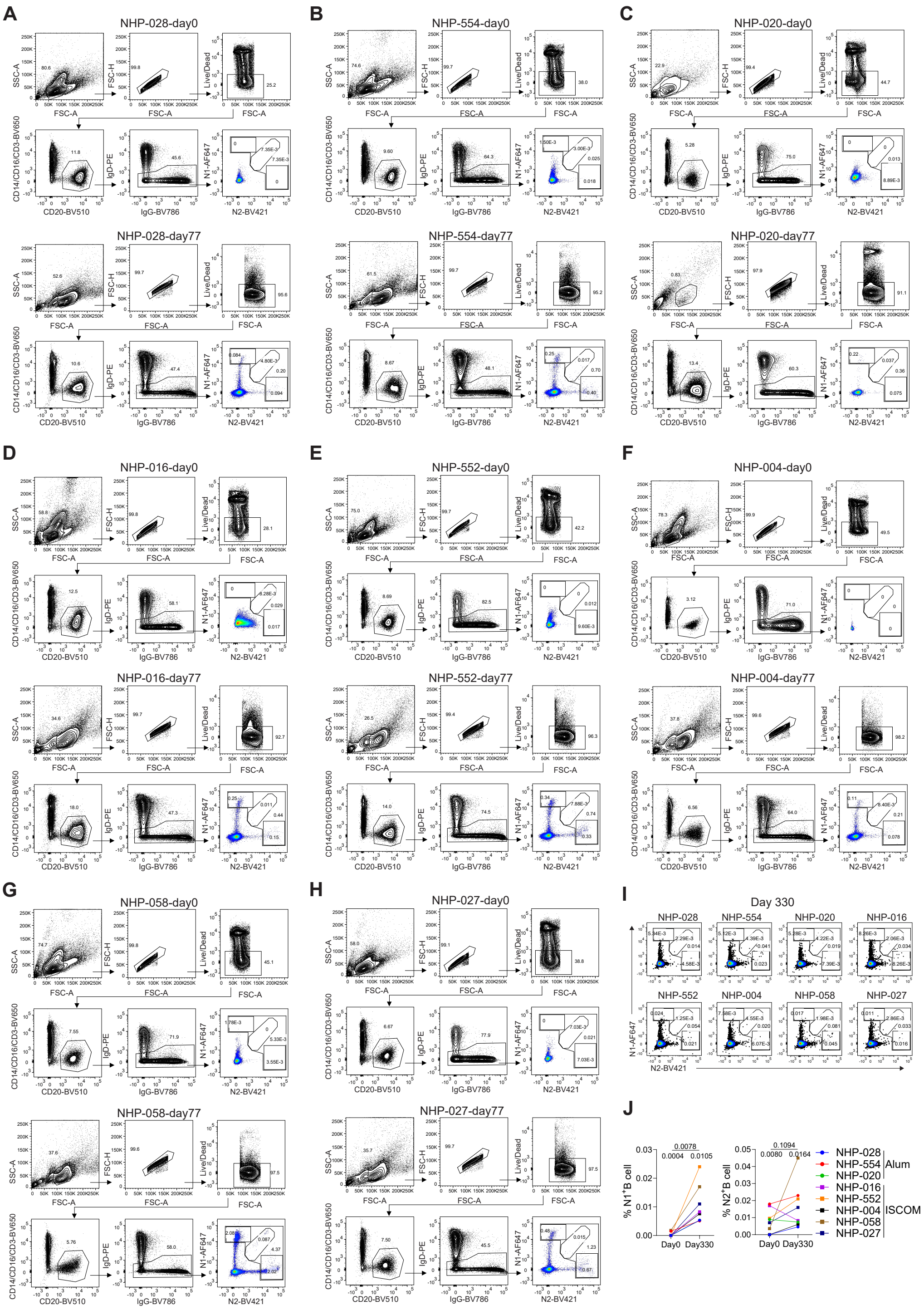


**Figure S5. Gating strategy for NA-specific MBCs in PBMCs from vaccinated NHPs. (A to H)** Gating strategy to identify NA-specific MBCs in each NHP at day 0 and day 77. **(I)** Representative flow plots show NA-specific MBCs in each NHP at day 330. **(J)** The frequencies of $N1^+$ or $N2^+$ MBCs in each NHP at day 0 and Day 330 were compared. The statistical significance was analyzed using the Wilcoxon matched-pairs signed-rank test, and *p*-values were indicated.

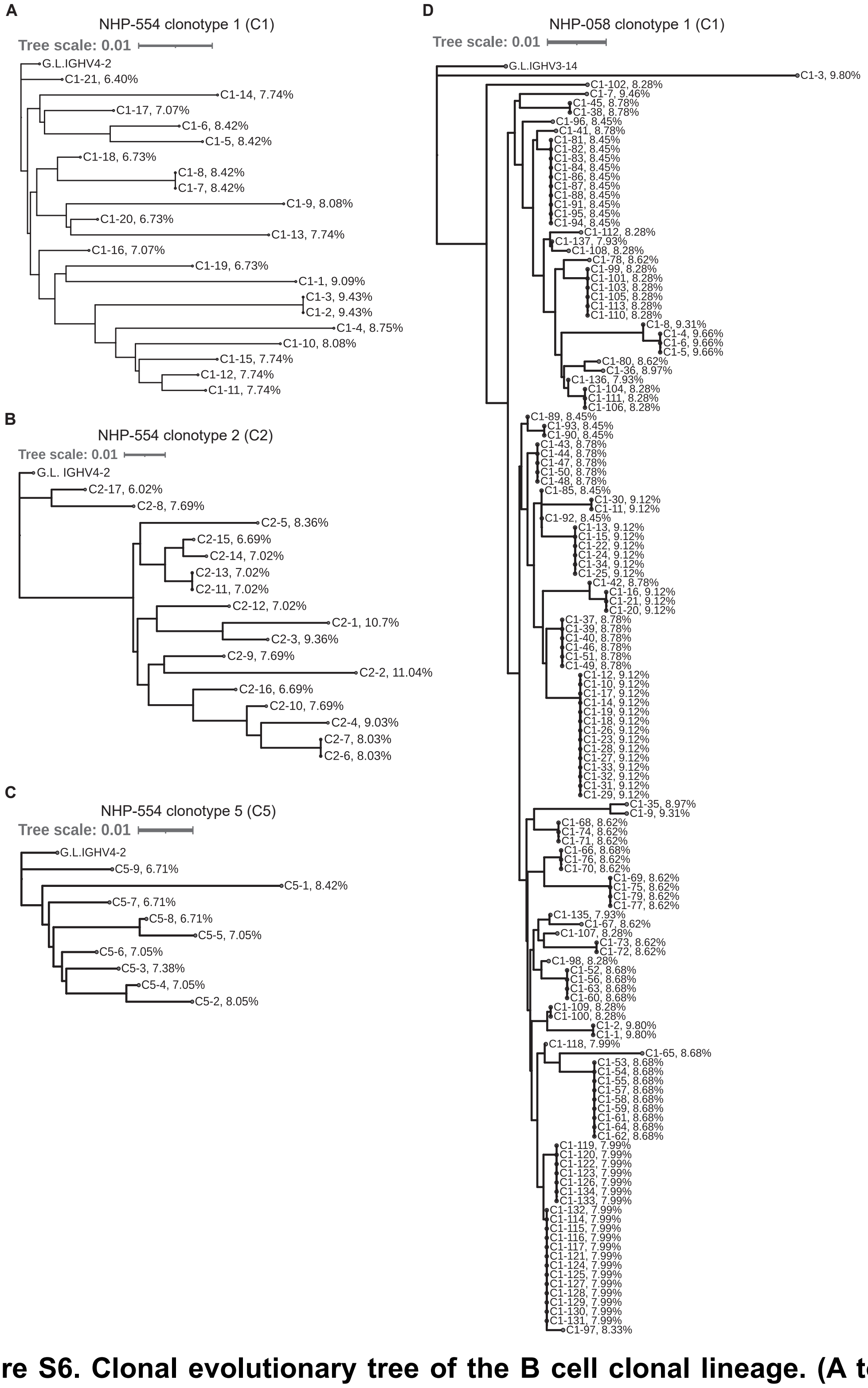


**Figure S6. Clonal evolutionary tree of the B cell clonal lineage. (A to D)** Clonal evolutionary trees based on heavy chain VDJ sequences, and the heavy chain *V* gene mutation rate are shown for NHP-554 clonotype 1 (C1) (A), clonotype 2 (C2) (B), clonotype 5 (C5) (C), and NHP-058 clonotype 1 (C1) (D) are shown, respectively. The scale bar represents a 0.01% change in nucleotides.

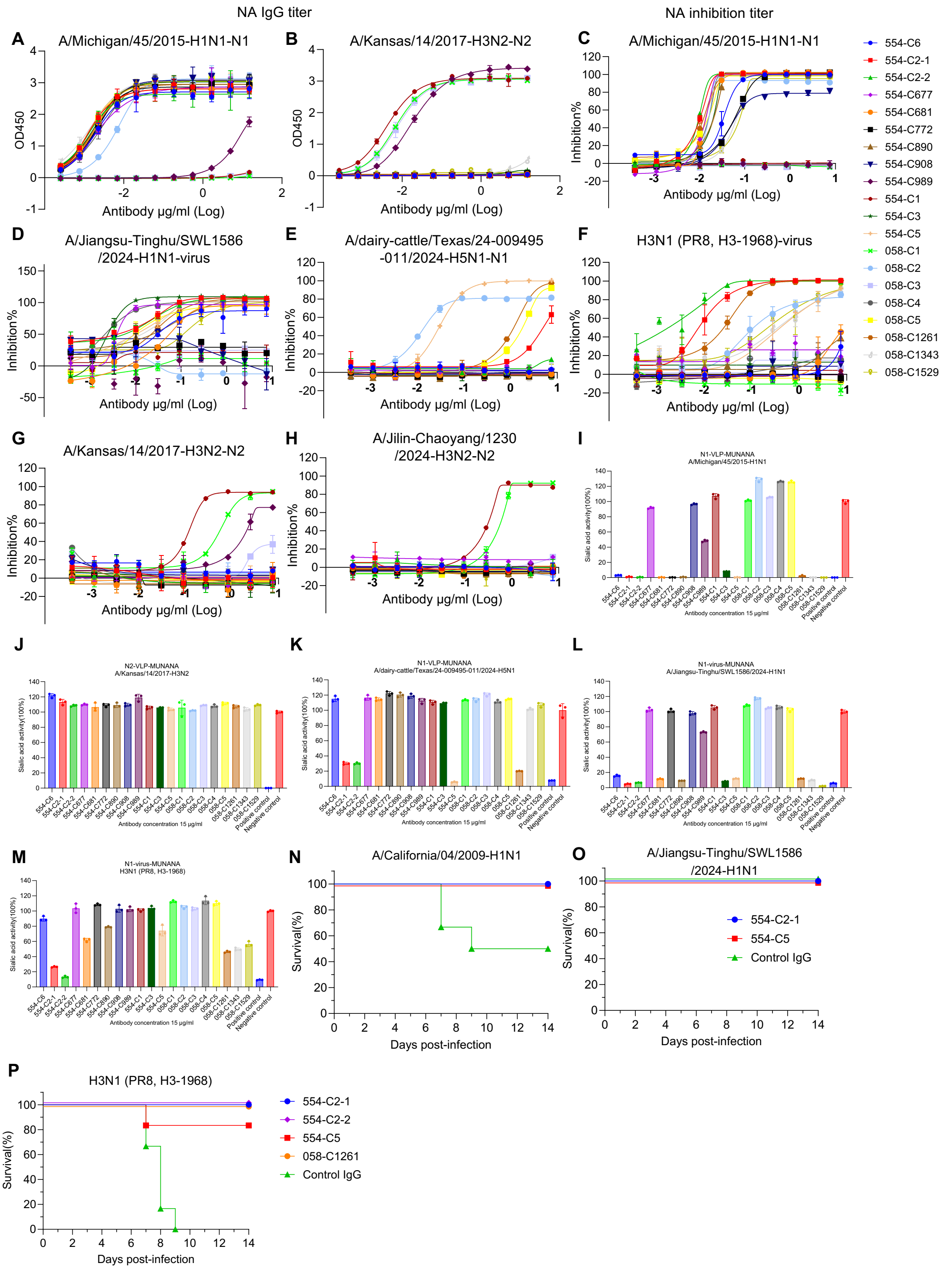


**Fig. S7. Binding, inhibitory, and *in vivo* protective activity of NA mAbs. (A and B)** The binding activity of mAbs to recombinant N1 (rN1, A/Michigan/45/2015-H1N1) and N2 (rN2, A/Kansas/14/2017-H3N2) was assessed by ELISA. **(C to H)** NI activities of mAbs against various influenza virus strains were determined by ELLA. The data are presented as mean ± s. d. The technical duplicates were utilized for each mAbs. **(I to M)** Inhibition of NA enzymatic activity by mAbs was assessed by MUNANA assay using indicated different influenza viruses or NA-VLPs. The technical triplicates were utilized for each mAbs. **(N to P)** The survival rates in BALB/c mice following prophylactic administration of the indicated NA mAbs and then challenging with the indicated influenza viruses. Mice (n = 6 to 7 per group) received an intraperitoneal injection of the indicated NA mAbs or an irrelevant human IgG (control IgG) 12 hours before being challenged with ~5 × $mLD_{50}$ of A/California/04/2009 (H1N1), 1 × $10^6$ PFU of A/Jiangsu-Tinghu/SWL1586/2024 (H1N1) or ~3 × $mLD_{50}$ of H3N1 (PR8, H3-1968), and the body weight were monitored during the flowing 14 days. The mice were euthanized when the reduction of body weight reached 25% of initial weight.

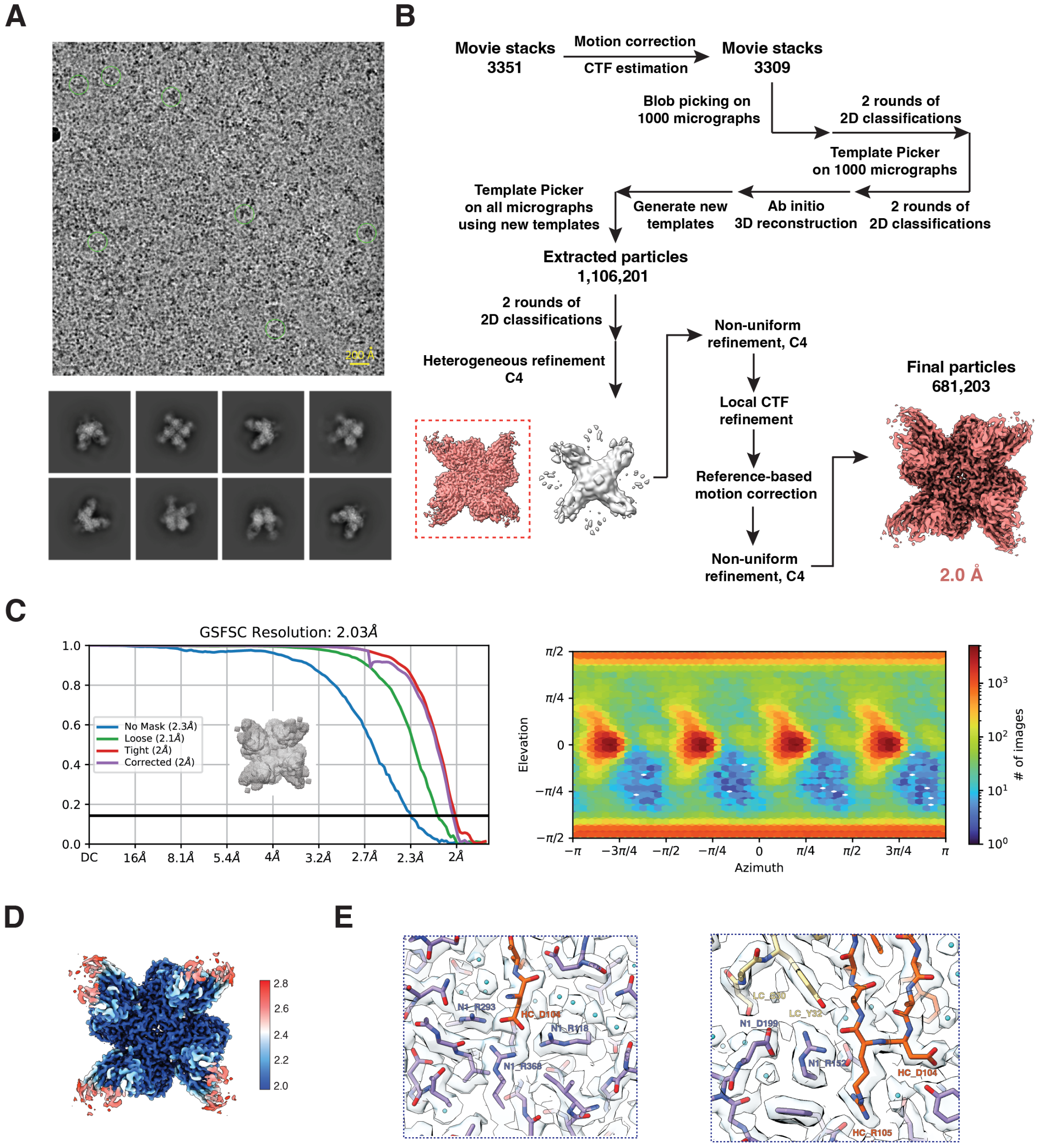


**Figure S8. Cryo-EM structure of 554-C2-1 with tetrameric N1. (A)** The top image shows the representative cryo-EM micrograph (defocus at around 2 μm) of the complex. The green circles, with a diameter of 200 Å, indicate representative particles picked for downstream analysis. The bottom images display some representative 2D class averages during data analysis. **(B)** Image processing workflow for the dataset. The class highlighted in color and boxed was used for further analysis. **(C)** The left panel is the Golden Standard Fourier Shell Correlation (GSFSC) curve for the 3D reconstruction. The black line indicates the 0.143 criterion. The mask used for refinement and resolution estimation is shown as a mesh. The right panel shows the Directional

Distribution of the 3D reconstruction. **(D)** The local resolution estimation of the cryo-EM map is shown. The final unsharpened map is colored based on local resolution estimation, and the scale bar represents values in angstroms (Å). **(E)** Cryo-EM densities with fitted model for the complex. The sharpened map, contoured at 0.15, was used. Models were shown as sticks with some key interacting residues labeled.

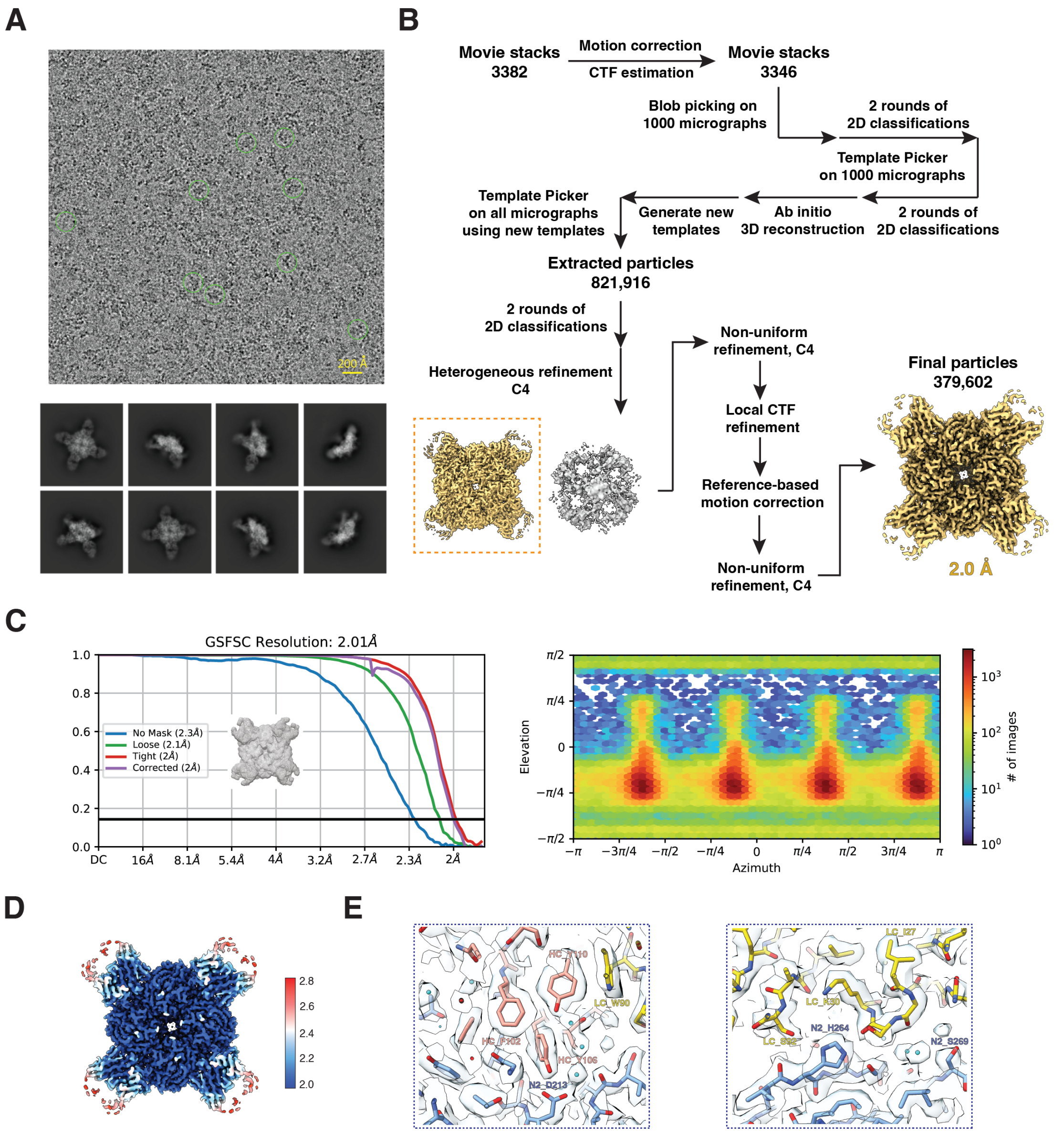


**Figure S9. Cryo-EM structure of 554-C1 with tetrameric N2. (A)** The top image shows the representative cryo-EM micrograph (defocus at around 2 μm) of the complex. The green circles, with a diameter of 200 Å, indicate representative particles picked for downstream analysis. The bottom images display some representative 2D class averages during data analysis. **(B)** Image processing workflow for the dataset. The class highlighted in color and boxed was used in the further analysis. **(C)** The left panel is the Golden Standard Fourier Shell Correlation (GSFSC) curve for the 3D reconstruction. The black line indicates the 0.143 criterion. The mask used for refinement and resolution estimation is shown as a mesh. The right panel shows the Directional Distribution of the 3D reconstruction. **(D)** The local resolution estimation of the cryo-EM map is shown. The final unsharpened map is colored based on local resolution estimation, and the scale bar represents values in angstroms (Å). **(E)** Cryo-EM densities with fitted model for the complex. The sharpened map, contoured at 0.15, was used. Models were shown as sticks with some key interacting residues labeled.

| | Heavy Chain | | | | | | Light Chain (Lambda/Kappa) | | | | |
|---|---|---|---|---|---|---|---|---|---|---|---|
| | V | D | J | V-mutation% | CDR3 | CDR3 length | V | J | V-mutation% | CDR3 | CDR3 |
| 554-C1-1 | IGHV4-2*01 | IGHD3-3*01 | IGHJ4*01 | 9.09 | CARSNFWNGYP**DR**YYFDYW | 19 | IGLV3-6*01 | IGLJ3*01 | 4.48 | CQVWDSSSKYALF | 13 |
| 554-C2-1 | IGHV4-2*01 | IGHD2-33*01 | IGHJ6*01 | 10.70 | CAREKWS**DR**LNYGLDSW | 17 | IGKV1S1*01 | IGKJ1*01 | 3.85 | CQQYDSLPRTF | 11 |
| 554-C2-2 | IGHV4-2*01 | IGHD1-32*01 | IGHJ6*01 | 11.04 | CAREKWT**DR**LNFGLDSW | 17 | IGKV1S1*01 | IGKJ1*01 | 5.94 | CQQYDSLPRTF | 11 |
| 554-C3 | IGHV3-7*01 | IGHD4-29*01 | IGHJ4*01 | 13.19 | CGKAPTNDYGTNW | 13 | IGLV3-1*01 | IGLJ2*01 | 7.64 | CQVWDSSNDHWVF | 13 |
| 554-C5 | IGHV4-2*01 | IGHD6-37*01 | IGHJ4*01 | 8.42 | CARPFRWYVDYW | 12 | IGKV1S22*01 | IGKJ4*01 | 7.39 | CQQHDSYPLTF | 11 |
| 554-C6 | IGHV3-13*01 | IGHD2-21*01 | IGHJ4*01 | 3.69 | CTRSHYCT**DR**GCYGTFDYW | 19 | IGKV1-9*01 | IGKJ2*01 | 2.82 | CLQYNSHPYNF | 11 |
| 554-C677 | IGHV3-7*01 | IGHD6-25*01 | IGHJ4*01 | 13.18 | CSGDGGS**DR**QMVFDSW | 16 | IGLV3-6*01 | IGLJ1*01 | 6.92 | CQVWDTSSKYDYIF | 14 |
| 554-C681 | IGHV4-2*01 | IGHD3-22*01 | IGHJ4*01 | 6.76 | CASWVWS**DR**REHFDSW | 16 | IGKV1-21*01 | IGKJ1*01 | 5.63 | CQQYSGYPWTF | 11 |
| 554-C772 | IGHV3-14*01 | IGHD2-33*01 | IGHJ4*01 | 10.81 | CAKDRWS**DR**NKGYFDYW | 17 | IGKV1-23*01 | IGKJ2*01 | 3.62 | CFQYNSYPYNF | 11 |
| 554-C890 | IGHV4-2*01 | IGHD6-37*01 | IGHJ6*01 | 7.02 | CARDAWT**DR**MRYNGFDSW | 18 | IGKV1S22*01 | IGKJ3*01 | 7.39 | CQQHDSYPFTF | 11 |
| 554-C908 | IGHV2-1*01 | IGHD3-22*01 | IGHJ3*01 | 3 | CARGSWSA**DR**FSAFDFW | 17 | IGLV1-14*01 | IGLJ1*01 | 4.08 | CQSYDSSLSANYIF | 14 |
| 554-C989 | IGHV3-9*01 | IGHD3-3*01 | IGHJ5-1*01 | 8.68 | CTTGGIFGVV**DR**WNNWFDVW | 20 | IGLV3-4*01 | IGLJ6*01 | 4.17 | CQVWDSTSDHDVF | 13 |
| 058-C1 | IGHV3-9*01 | IGHD5-12*01 | IGHJ4*01 | 9.80 | CAKDSGYTYTYLDHW | 15 | IGKV1-15*01 | IGKJ1*01 | 2.79 | CQQGNSNPPWTF | 12 |
| 058-C2 | IGHV4-2*01 | IGHD3-3*01 | IGHJ4*01 | 11.11 | CARSNNLWTGFYYFDFW | 17 | IGLV1S1*01 | IGLJ3*01 | 1.36 | CQSYDSSLSGYVLF | 14 |
| 058-C3 | IGHV3-6*01 | IGHD3-28*01 | IGHJ6*01 | 9.03 | CYSYTGYSYYALDSW | 15 | IGKV1-9*01 | IGKJ1*01 | 4.55 | CLQYKRDPRTF | 11 |
| 058-C4 | IGHV4-2*01 | IGHD2-27*01 | IGHJ4*01 | 9.60 | CARDNNCRDTYCYAGVFVYW | 20 | IGLV3-1*01 | IGLJ2*01 | 9.38 | CQVWDNSRDHGLF | 13 |
| 058-C5 | IGHV3-14*01 | IGHD3-28*01 | IGHJ4*01 | 9.83 | CARGWIDDYGSGYYTFDYW | 19 | IGKV3S7*01 | IGKJ3*01 | 4.61 | CQQYNNWNTF | 10 |
| 058-C1261 | IGHV4-2*01 | IGHD2-33*01 | IGHJ5-1*01 | 10.14 | CVRAPWS**DR**LFSGWFDVW | 18 | IGKV3S9*01 | IGKJ1*01 | 3.90 | CQQYNDWDRTF | 11 |
| 058-C1343 | IGHV4-2*01 | IGHD3-28*01 | IGHJ1*01 | 7.46 | CARGLYYD**DR**GPPRAPEFW | 19 | IGKV1-13*01 | IGKJ2*01 | 2.11 | CLQGYTTPYSF | 11 |
| 058-C1529 | IGHV3-14*01 | IGHD3-28*01 | IGHJ4*01 | 14.92 | CARARLLYYY**DR**DTNVRKRYYFDYW | 25 | IGLV6-2*01 | IGLJ6*01 | 1.35 | CQSADDSYNEVF | 12 |

**Table S1. The VDJ usage, CDR3 lengths, and V gene mutation rate of heavy and light chains of selected mAbs from NHPs 058 and 554 are shown.**

| Structure | 554-C2-1-N1 | 554-C1-N2 |
|---|---|---|
| **Data Accession** | | |
| PDB | 21TM | 21TN |
| EMDB | EMD-67987 | EMD-67988 |
| **Data Collection** | | |
| Microscope | Titan Krios G4 | |
| Voltage (kV) | 300 | |
| Exposure navigation | Stage Movement/Image Shift | |
| Automation software | EPU | |
| Detector | Falcon 4i | |
| Energy filter | 10 | |
| Nominal magnification | 130 k | |
| Pixel Size (Å) | 0.92 | |
| Electron exposure ($e^-$/Å$^2$) | 50 | |
| Defocus range (μm) | -1.1 to -2.2 | |
| Micrographs collected | 3351 | 3382 |
| **Reconstruction** | | |
| Software | CryoSPARC v4 | |
| Micrographs used | 3309 | 3346 |
| Particles extracted | 1,106,201 | 821,916 |
| Particles used in refinement | 681,203 | 379,602 |
| Symmetry | C4 | |
| Overall resolution (Å) FSC=0.143 (masked) | 2.0 | 2.0 |
| Map sharpening B-factor (Å$^2$) | -50.0 | -50.0 |
| **Model Refinement** | | |
| Software | Phenix | |
| Model Composition | | |
| Non-hydrogen atoms | 19816 | 20164 |
| Protein residues | 2452 | 2456 |
| Water | 784 | 692 |
| Ligands | 16 | 40 |
| B factors (Å$^2$) | | |
| Protein | 24.71 | 21.15 |
| Water | 23.75 | 17.62 |
| Ligands | 36.33 | 29.72 |
| R.M.S. deviations | | |
| Bond length (Å) | 0.006 | 0.005 |
| Bond angle (˚) | 1.123 | 1.111 |
| Ramachandran statistics (%) | | |
| Outliers | 0.00 | 0.00 |
| Allowed | 2.84 | 2.71 |
| Favored | 97.16 | 97.29 |
| MolProbity score | 1.22 | 1.39 |
| All-atom clashscore | 2.83 | 4.92 |
| Poor rotamers (%) | 0.52 | 0.38 |
| Model vs. Map FSC FSC=0.5 (masked, Å) | 2.1 | 2.1 |

**Table S2. Statistics for Cryo-EM data collection and refinement.**
